# Zapit: Open Source Laser-Scanning Photostimulation For Neuroscience

**DOI:** 10.1101/2024.02.12.579892

**Authors:** Michael Lohse, Oliver M Gauld, Maja T Quigley, Ainiah Masood, Chaofei Bao, Jingjie Li, Nathaniel J Miska, Gerion Nabbefeld, Quentin Pajot-Moric, Simon Townsend, Peter Vincent, Angela Y Yang, Nikolaos Zervogiannis, Athena Akrami, Chunyu A Duan, Jeffrey C Erlich, Sonja B Hofer, Thomas D Mrsic-Flogel, Philip Coen, Robert AA Campbell

## Abstract

Optogenetic tools are indispensable for understanding the causal neural mechanisms underlying animal behavior, providing millisecond-precision control over genetically defined neural populations. Random-access laser-scanning optogenetics is a powerful and underutilized method for transiently manipulating cortical activity in mice. Despite the utility of this technique, no general-purpose open-source implementation currently exists, and potential users need expertise across optics, real-time hardware control, and programming. This represents a major barrier to adoption, particularly for newly established research groups. Here we present ‘Zapit’, the first open-source general-purpose platform for random-access laser-scanning optogenetic experiments in head-fixed mice. Zapit is fully documented, has a user-friendly GUI, works in stereotaxic coordinates, and comes with easy to build hardware options that extend functionality beyond any published system. We validate Zapit’s performance through electrophysiological recordings and cortical photoinhibition in behaving mice. Zapit is a novel and innovative tool with the potential to democratize laser-scanning optogenetics and significantly increase uptake throughout the scientific community.

## 1 Introduction

Optogenetics is now the principal experimental tool for manipulating neural activity in systems neuroscience due to its high temporal precision, non-invasive options, reversibility, and genetically and spatially restricted specificity (Boyden *et al*. 2005; O’Connor, Huber, and Svoboda 2009; Häusser 2014; Li *et al*. 2019; Emiliani *et al*. 2022). Optical access to the brain to stimulate opsins is predominantly achieved through chronically implanted fibers (e.g. Mayrhofer *et al*. 2019; Babl, Rummell, and Sigurdsson 2019; Lohse *et al*. 2021; Duan *et al*. 2021; Emiliani *et al*. 2022), or implanting a window over the cortical site (e.g. Schneider, Sundararajan, and Mooney 2018; Liu, Huberman, and Scanziani 2016). Implanted fibers allow for targeted light delivery in both head-fixed and freely moving behavioral paradigms and enable access to deep brain areas. However, the physical footprint and chronic nature of implants can restrict manipulation to a predetermined and limited number of sites.

In mice, the skull is thin enough (~250 *µm*; Ghanbari *et al*. 2019) to be remarkably transparent (Steinzeig, Molotkov, and Castrén 2017). This transcranial optical access allows for non-invasive, laser-scanning (random access) optogenetic photostimulation across the entire dorsal cortical surface. Various approaches exist to achieve this goal. An optical fiber can be mounted on a motorized micro-drive (Zatka-Haas *et al*. 2021), a digital micro-mirror device (DMD, Allen 2017) can project arbitrary illumination patterns (e.g. Chong *et al*. 2020), or a pair of galvanometric scan mirrors (galvos) can direct a focused laser beam to specified positions on the skull (Guo, Li, *et al*. 2014; Li *et al*. 2016; Heindorf, Arber, and Keller 2018; Inagaki *et al*. 2018; Pinto *et al*. 2019; Keller, Roth, and Scanziani 2020; Esmaeili *et al*. 2021; Voitov and Mrsic-Flogel 2022; Pinto, Tank, and Brody 2022; Coen *et al*. 2023; Gauld *et al*. 2025). A galvanometric approach to laser targeting offers the best utility-to-affordability trade-off: scanners are much faster than motorized drives, and because galvo mirrors effectively direct all light to the target site, the required laser power is much lower than a DMD. The high speed of the scanners nonetheless allows the experimenter to silence multiple locations near simultaneously.

While laser-scanning optogenetics approaches have been established in the field for several years, their use remains limited to a small number of technically sophisticated research groups. As these research groups have focused their publications on novel experimental results, it remains impractical for other groups to reproduce these methods without pre-existing expertise spanning optics, electrical engineering, and hardware control (Fig. S1). This represents a significant barrier, especially for newly established research groups. This is further compounded if the experimental aims differ from the original publication, or if specific parts are unavailable or custom-made. Therefore, despite its utility and potential, dissemination of laser-scanning photostimulation technology throughout the neuroscientific community has been limited.

Community-targeted versions of intricate experimental techniques facilitate widespread adoption: recent examples include comprehensive protocols and software tools for widefield calcium imaging (Couto *et al*. 2021) and two-photon all-optical interrogation (Russell *et al*. 2022). Furthermore, the widespread success of open-source hardware tools such as ‘Open-Ephys’ (Siegle *et al*. 2017) and the ‘Pulse-Pal’ pulse generator (Sanders and Kepecs 2014) demonstrate how accessible open-source versions of existing approaches can have a transformative effect on experimental neuroscience.

Here we present ‘Zapit’, an open-source galvo-based photostimulation system. Zapit integrates both hardware and software into a user-friendly experimental tool with functionality beyond that of any published system. It features a low-cost design based on commercially available components, with step-by-step build, usage, and troubleshooting instructions to promote accessibility. The modular hardware design allows users to modify the system to their experimental needs, including adaptations that support several stimulation wavelengths. The accompanying software facilitates semi-automated system calibration, flexible stimulus design and delivery, and is maintained by a community of developers and experimentalists. The system can interface with a range of programming languages for further integration into existing behavioural and experimental protocols. Through electrophysiological recordings, we confirm that Zapit’s flexible and accessible design matches the spatiotemporal resolution of existing laser-scanning solutions (Pinto, Tank, and Brody 2022, Guo, Li, *et al*. 2014, Li *et al*. 2019). Finally, we present proof-of-principle experiments demonstrating that Zapit can evoke robust effects across different sensory and motor systems in awake behaving mice, highlighting the utility and effectiveness of the system in revealing causal links between cortical areas and behavior.

## 2 Results

## 2.1 Operating principle

Zapit is a combined hardware and software solution for galvo-based photostimulation of the mouse dorsal cortex. We implemented a hardware design similar to Pinto *et al*. (2019) and Pinto, Tank, and Brody (2022), where a laser is directed into an XY galvo scanner, targeted to the animal via a dichroic mirror, and focused onto the sample with a scan lens. The scan lens doubles as an objective, imaging the sample and any excited auto-fluorescence onto a camera via a tube lens (Figure 1A). The camera provides a real-time image of the sample under the objective. This low-cost, accessible design is composed of commercially available parts (primarily Thorlabs) for easy set-up and usage (Figure 1B). The galvo waveforms are shaped to minimize mechanical noise from mirror movement, which should be consistent across trials. However, for use-cases where sound isolation is critical, we provide design-files for a custom enclosure for additional sound attenuation.

**Figure 1.**
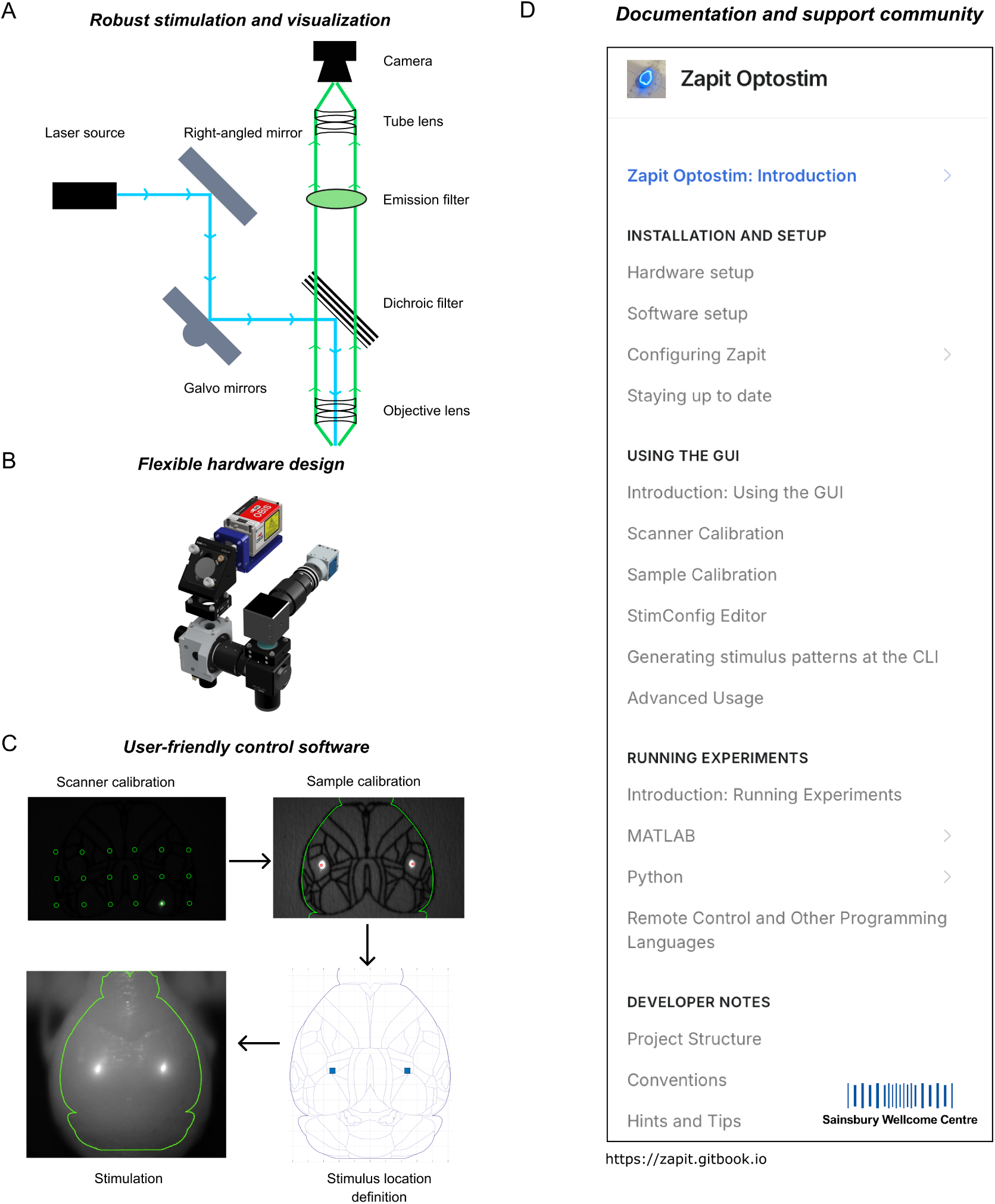
Zapit system overview. **A.** System schematic. The unexpanded laser beam (cyan) is directed onto the galvos and then targeted to the sample via a dichroic mirror. A lens focuses the laser beam onto the sample. The sample is imaged onto a camera (green) using this same ‘scan lens’ as an objective, combined with a tube lens in front of the camera. **B.** Our Zapit design is low-cost and largely consists of commercially available parts (custom parts shown in purple). This design can be modified to meet user needs, such as component orientation. A full parts list is available on Github. **C.** User-friendly control software. The software guides the user through sequential scanner and sample calibration steps (top left and right, respectively). From here, the user can define stimulus locations (bottom right), to then deliver stimulation (bottom left) to said locations. The end result is that calibrated stimuli are projected onto the imaged brain surface for real-time visualization. **D.** Full usage documentation and troubleshoot support. A dedicated website (zapit.gitbook.io) is available for setting up Zapit hardware and software, usage instructions and troubleshooting steps. Further support is available from the creators and wider community for user-specific implementation needs and issues.

The Zapit software comprises a MATLAB-based, Windows-run, user-friendly GUI to facilitate calibration and experimental integration. Calibration takes ~1 minute and involves two steps (Figure 1C): **1** mapping scanner positions to pixel-space in the camera image, and **2** mapping stereotaxic coordinates onto the image of the exposed skull. Once calibrated, the beam can be targeted via stereotaxic coordinates, and a simple API allows the user to integrate stimulation into an experimental paradigm. The software supports stimulation of multiple locations in a given experimental trial, with each location stimulated sequentially at a specified frequency (40 Hz in experiments described here). The laser intensity ramps down at trial end to limit rebound effects (Li *et al*. 2019). Detailed build instructions, operating guidelines and support are available via our GitHub repository (Figure 1D).

### 2.2 Hardware Design

Zapit is designed with commercially available components to ease construction and maximize affordability (Figure 2). The system is built around ThorLabs GVS002 scanners which direct the beam via an NI DAQ, and images are acquired using a USB-3 Basler acA1920-40um camera. The beam feeds into the galvo scan head via an adjustable fold-mirror. A second, fixed, fold-mirror placed before the camera keeps the system compact. To focus and image the beam we used an f=200 mm Plössl objective lens composed of two ThorLabs AC254-400-A (f=400 mm) achromats. A Plössl is a compound lens composed of two identical achromatic doublets arranged such that their positive (convex) surfaces are near touching. This arrangement has substantially reduced optical aberrations and distortions compared with a single achromatic doublet of equivalent focal length. We used an f=100 mm Plössl tube lens (2x ThorLabs AC254-200-A achromats), resulting in a magnification of 0.5x which comfortably fits the mouse brain onto the camera sensor.

**Figure 2.**
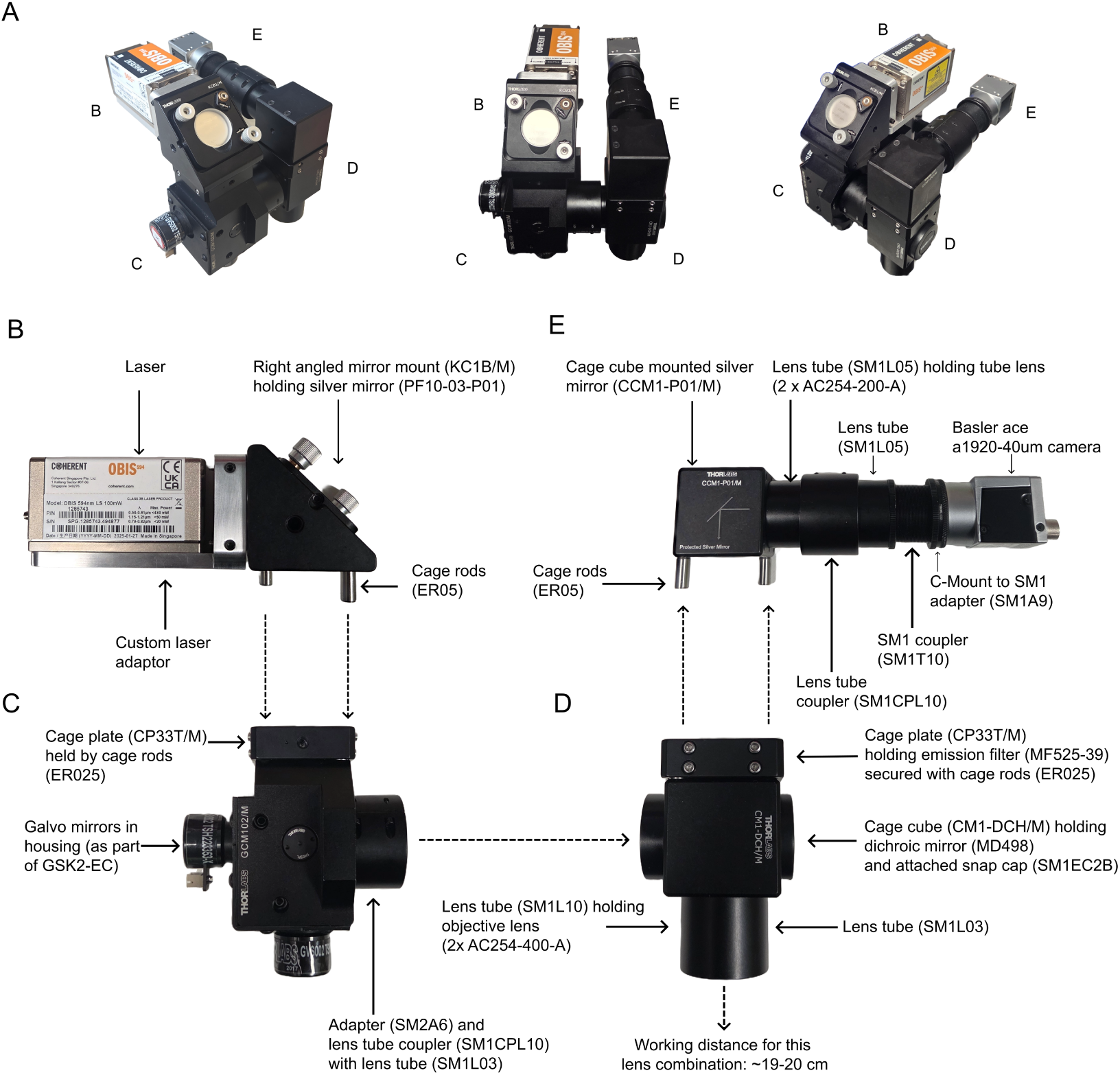
Hardware components of Zapit. All parts, except for the laser and laser adaptor, are available from Thorlabs, with appropriate part numbers. Dashed arrows indicate where sections should be joined. A comprehensive parts list and assembly instructions with visual aids are available on Github. Readers unfamiliar with the basics of optical scanning can get more information from Schottdorf *et al*. 2025. **A**. Left side, middle and right side views of a fully-constructed Zapit system. Smaller letters correspond to individual sections below. **B**. Laser attached to custom adapter connected to associated right angled-mirror. **C**. Galvo mirrors with attached connector parts. **D**. Dichroic filter in cage cube with objective lens in lens tube and emission filter in cage plate. **E**. Camera attached to connecting tubes, with one lens tube containing tube lens.

A variety of lasers, dichroics, and emission filters can be chosen to support diverse applications with different opsins. A complete parts list, with suggested alternatives, and assembly instructions, are available on GitHub. For a setup with a 473 nm laser (we used a Coherent Obis) targeting Channelrhodopsin-2 (ChR2), we recommend a ThorLabs MD498 dichroic to direct the light to the sample and allow reflected light to reach the camera. An MF525-39 emission filter in front of the camera allows visualization of autofluorescence from the target location without contamination from the excitation laser. We modelled a Zapit variant with an f=150 mm objective in Zemax and calculated the theoretical laser spot as having a PSF with a FWHM of 74 µm. We measured the true PSF full width half max as 91 µm (Supplemental Fig. S2). Spot size scales linearly with focal length and so the f=200 mm lens described above would have a PSF of ~120 µm. Nonetheless, resolution is limited by scattering which creates a much larger spot size within the brain (~1 mm radius, (Li *et al*. 2019)). Since the beam diameter entering the scan lens is ~0.8 mm, an f=200 mm objective will yield an excitation NA of about 0.002, meaning the beam is always in focus over the curved skull. The Rayleigh range (*Z_R_*) defines the axial distance from focus where the beam diameter doubles; in our system *Z_R_* = 37.6 *mm*.

The modular and open-source nature of Zapit means that new configurations can be constructed, such as employing two lasers in one experiment to stimulate different opsins within one mouse (Supplemental Fig. S3). Future developments to Zapit’s control code will facilitate this use-case.

### 2.3 Software Design: calibration

The main Zapit GUI (Figure 3A-i) is launched by running start_zapit at the MATLAB command line. Using this GUI, calibration is a simple two-step procedure. First, we estimate the mapping from the voltage commands applied to the galvos, and the resulting laser location in the pixel-space of the camera image. This is achieved by scanning the beam over a grid of points and then conducting an affine transform between the intended and observed beam locations (Figure 3A-iii). Beam locations on the sample are automatically detected by the camera through image thresholding. The entire scanner calibration is therefore an automated procedure, initiated by a single button-click in the GUI. Following scanner calibration, the user can manually trigger ‘on the fly’ targeted photostimulation by clicking on locations the sample image in Point Mode (Figure 3A-iv).

**Figure 3.**
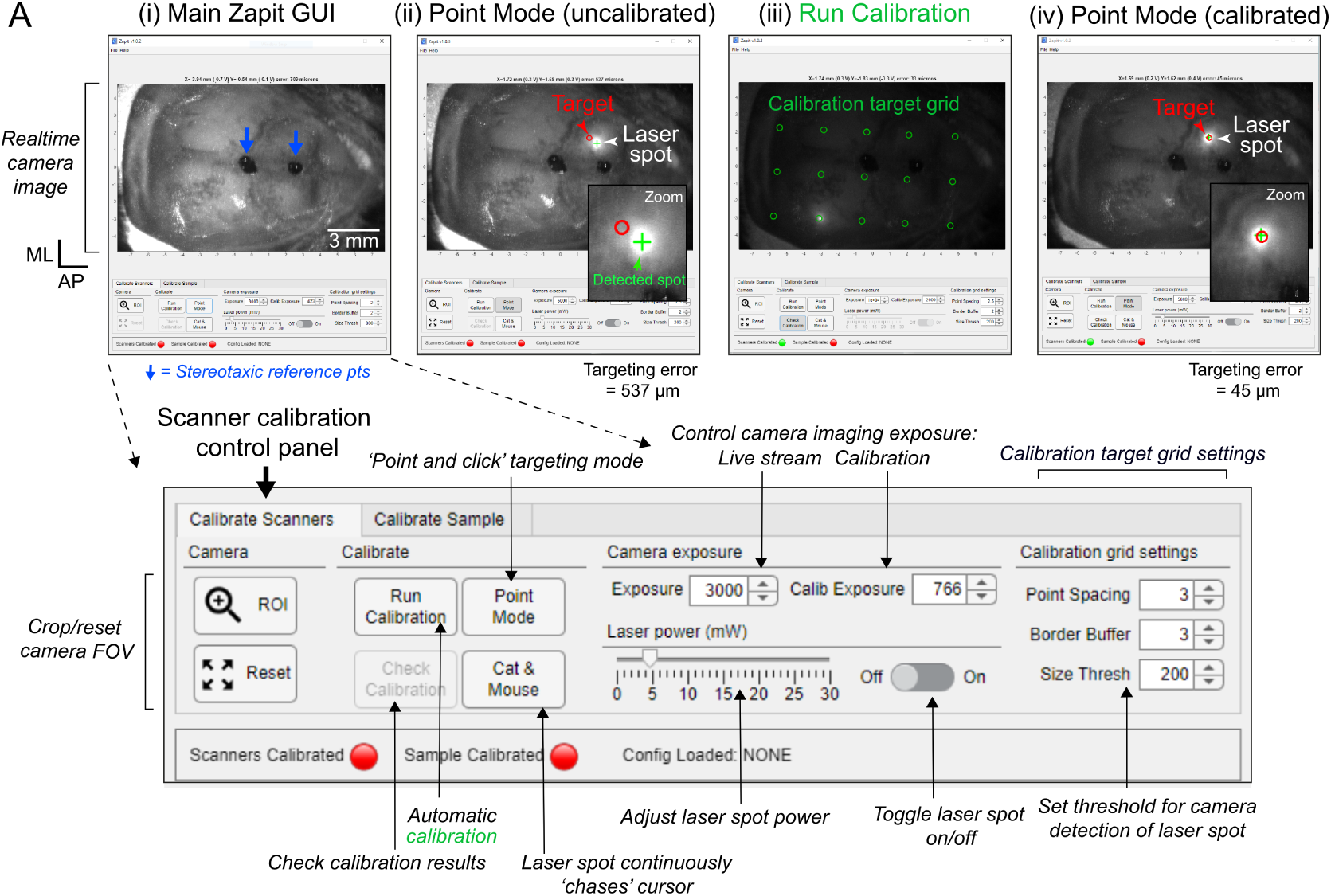
Scanner calibration using the Zapit GUI. **A**. (i) The Zapit GUI displays a live image feed from the camera. This example shows the dorsal surface of the skull with the two blue arrows indicating the location of bregma and a position 3 mm anterior from bregma. The panel below shows an close-up view of the GUI control panel, highlighting options for calibration, controlling laser stimulation and camera imaging. (ii) In ‘Point Mode’, the user can manually click locations in the image FoV, and Zapit will target light to this location. However, when ‘Uncalibrated’, the actual beam location (bright point) does not match with the desired target location (red circle). The green ‘+’ indicates the automatic detection of the beam location on the sample. (iii) ‘Run Calibration’ initiates an automatic calibration routine to correct this targeting error. Zapit systematically moves the beam over a grid of points and conducts an affine transform between the camera-measured and intended beam positions. (iv) Once calibrated, the user can click on locations in the image and the beam goes to the intended target position with minimal error.

Following scanner calibration, the user calibrates the sample to brain atlas space by defining the position of the skull in the FoV using two landmarks, such as bregma and bregma +3 mm AP. These two landmarks are marked onto the skull during surgery (Supplemental Fig. S4). Whilst clicking these coordinates, a brain outline is dynamically positioned, scaled, and rotated, providing instant feedback on the calibration (Figure 4A-i-iii). This process assumes the skull is level under the objective, which should be considered during headplate implantation (Supplemental Fig. S4, and Methods). Once stimulus calibration is complete, photostimulation target locations can be specified in stereotaxic coordinates, relative to bregma (Figure 4A-iv).

**Figure 4.**
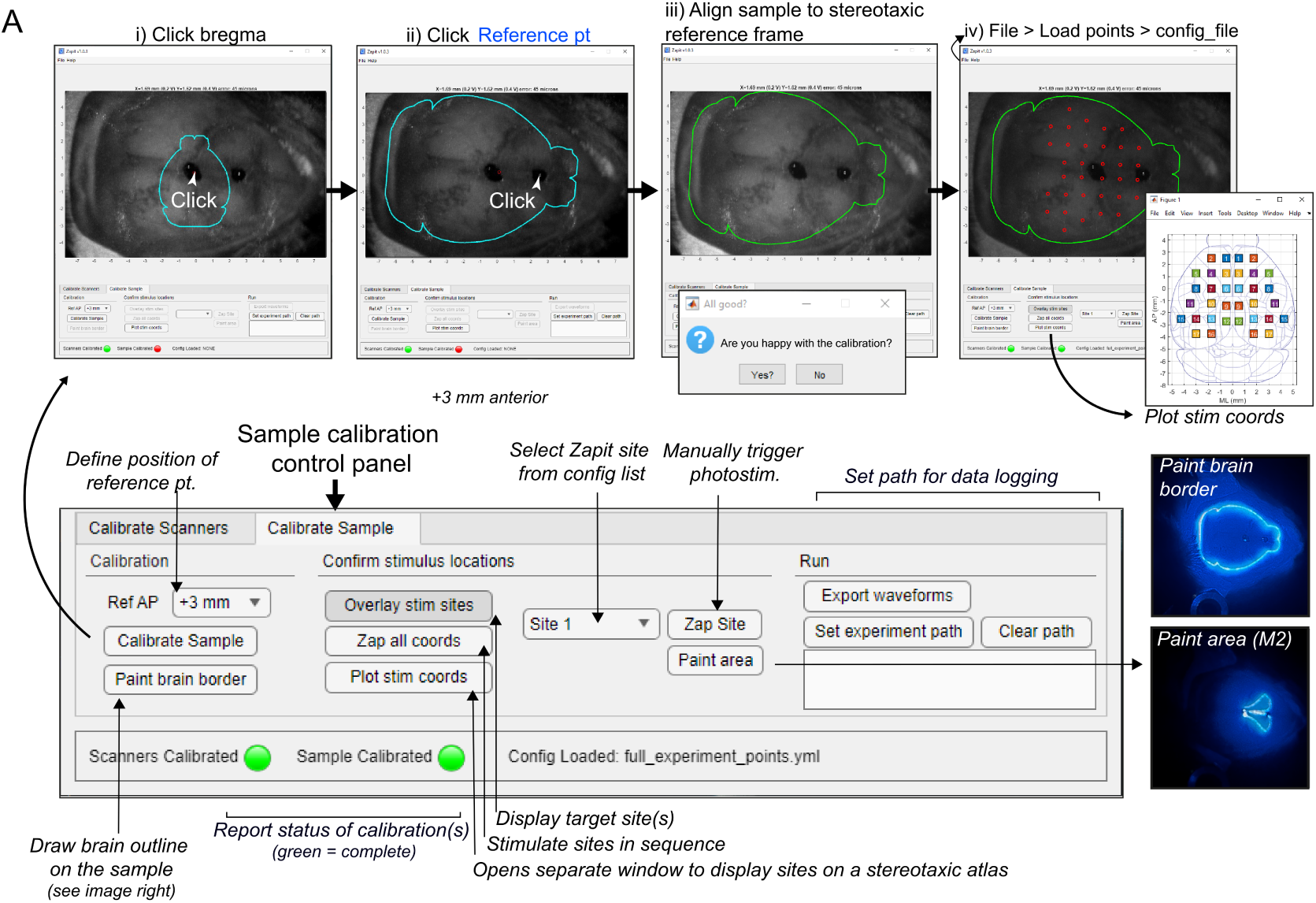
Sample calibration using the Zapit GUI. **A**. (i-ii) Sample calibration. The user defines the position of the skull in the FoV using two landmarks, such as bregma and bregma +3 mm. These coordinates can be marked onto the skull during a previous surgery using a stereotaxic frame. Whilst clicking these coordinates, a brain outline (blue) is dynamically positioned, scaled, and rotated, providing instant feedback on the calibration. (iii) When both reference points have been selected, the user will be prompted to confirm the calibration. Once confirmed, the brain outline will turn green. (iv) The user can then load in target points defined in coordinate space, and confirm correct targeting by stimulating the target locations (‘Zap Site’). There are additional scanning options that allow for the outline of the dorsal brain mask, or specific cortical area masks, to be continuously traced by the laser on the sample (as shown by the two example photos). These can be useful for visual alignment checks.

### 2.4 Software Design: defining stimulus locations

Stimulation target locations are defined in stereotaxic space and stored in a human-readable stimulus configuration file, which can be generated using the GUI (Figure 5A). This coordinate space is derived from the Allen Atlas zeroed to bregma (with bregma estimated at 5.4 mm AP in native space, as in Birman *et al*. 2023). Figure 5B shows an example of a finished stimulus set that includes unilateral, bilateral and multi-site stimulation conditions.

**Figure 5.**
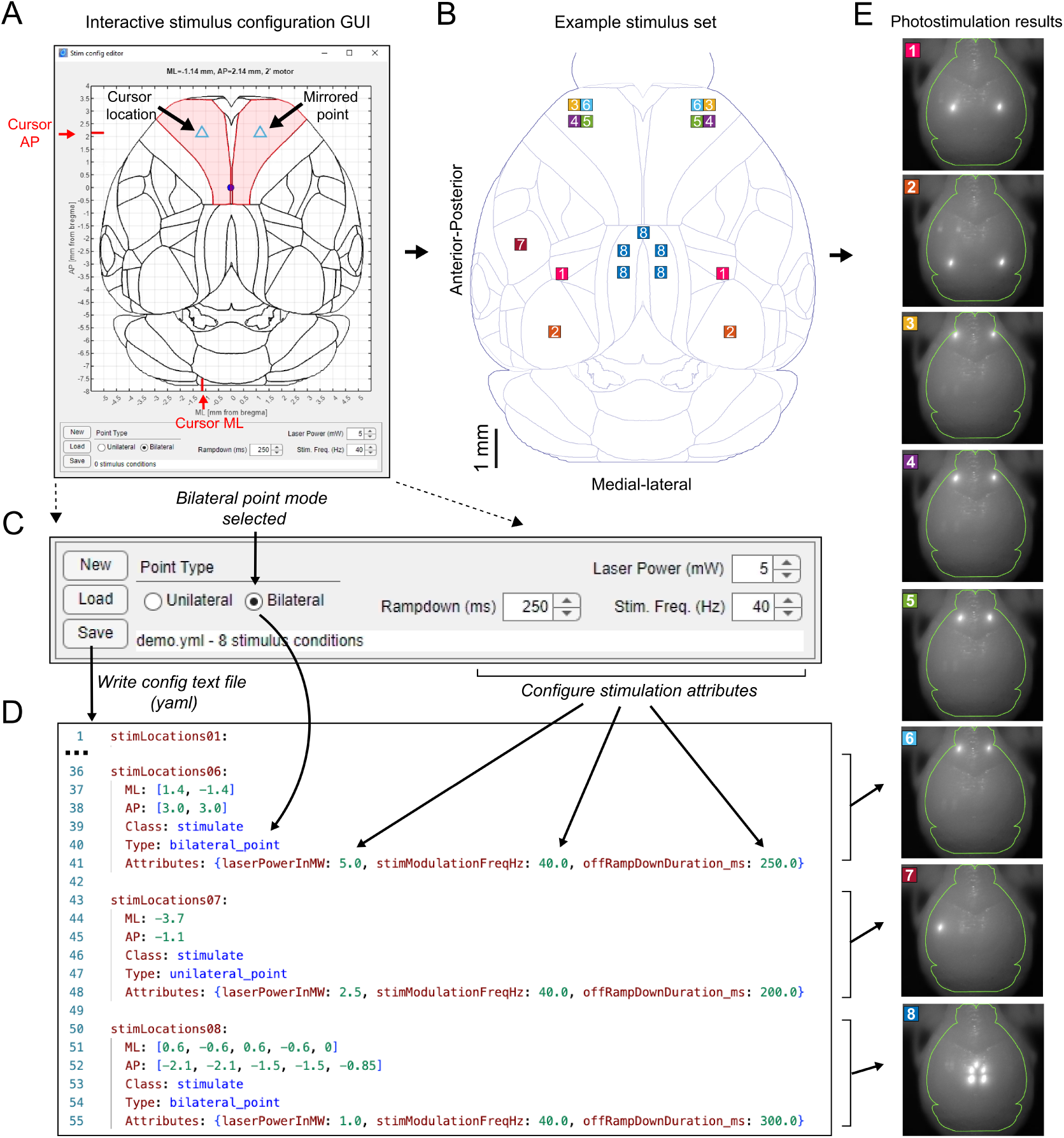
Generating photostimulation conditions using the interactive stimulus editor. **A**. Screenshot of the stimulus config editor. The user adds or modifies stimulus locations by clicking on the top-down view of the brain. Points can be added either freely (‘unilateral’ mode) or symmetrically on the left and right sides (‘bilateral’ mode). The laser power, stimulation frequency, and ramp-down time are set to default values for all stimuli using this GUI. The stimulus set is saved as a human-readable text file. Laser power, ramp-down, and stimulation frequency can be altered on a trial by trial basis by editing this file. **B**. A stimulus set superimposed onto the dorsal brain surface. All squares with the same number are associated with a single trial and will be stimulated together. **C**. Close-up of the GUI control panel showing stimulation parameters: laser power, rampdown duration, and stimulation frequency. The stimulus set can be saved and loaded as a human-readable text file. **D**. The corresponding stimulus configuration file (YAML) from the stimulus-set in B. Note that the file is abbreviated to highlight conditions 6-8 in the set. **E**. Results when targeting the individual trial patterns shown in panel B. Note these images show stimulation results on a 3D-printed mouse brain. The green brain outline shows the sample alignment result in the Zapit calibration GUI.

The stimulus configuration GUI allows the user to define general stimulation parameters in addition to stimulus locations, including ‘Laser Power’, ‘Rampdown’ and ‘Stimulation Frequency’ (Figure 5C). Note that the ‘Laser Power’ setting defines the *time-averaged power* in mW at the sample surface used for stimulation. In other words, if two points are being stimulated in a single trial and requested power is 2 mW then the laser would deliver 4 mW at each point on the sample with a 50% duty cycle. A power meter receiving light from just one of those points would report an average power of 2 mW.

Stimulus sets can also be generated and modified by directly editing the configuration text file (YAML; Figure 5D). The configuration file can be loaded via the main Zapit GUI enabling photostimulation of the defined target areas (Figure 5E).

### 2.5 Running Experiments

Once the system is calibrated and a stimulus configuration file is loaded, Zapit’s MATLAB API can be used to integrate photostimulation into the user’s experimental protocol. Precisely timed stimulation events are delivered in a trial-based paradigm using the sendSamples function (Figure 6A-B), which operates via the NI Data Acquisition (DAQ) card connected to the control PC (Figure 6C). Analog outputs AO0 and AO1 drive the galvo mirrors, AO2 controls laser power, and AO3 drives the masking light. Trials can be optionally hardware-triggered via PFI0, and the system can be controlled remotely from MATLAB, Python, or Bonsai using Zapit’s TCP/IP protocol (Figure 6B). Photostimulation ends with a gradual ramp-down in power (by default over a 250 ms time period) (Figure 6D). This limits large rebounds in activity, which are common following photoinhibition cessation with light-sensitive opsins (Li *et al*. 2019).

**Figure 6.**
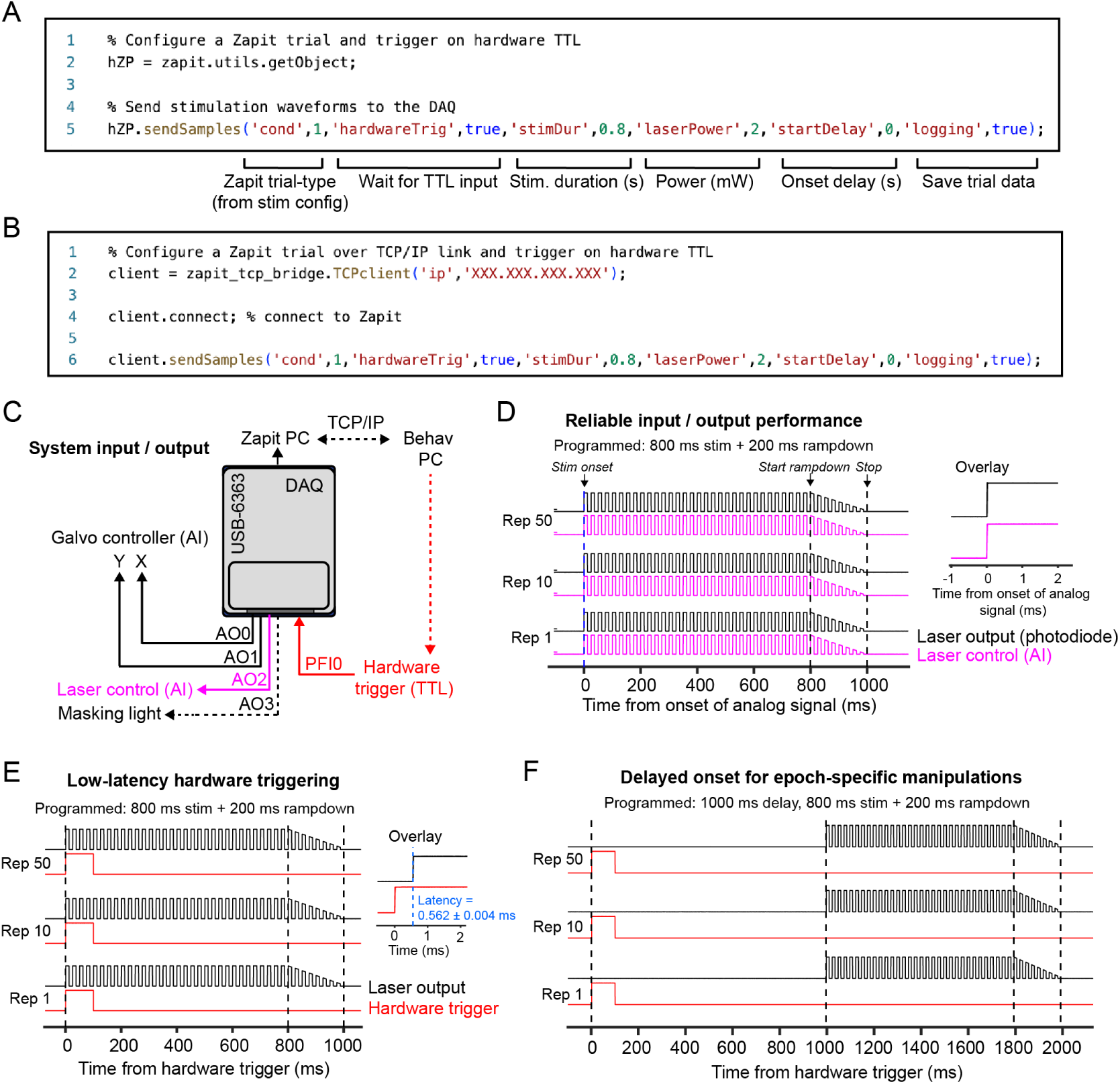
Reliable and precise photostimulation. **A.**MATLAB code snippet to run a stimulation paradigm with TTL input, with users being able to adjust trial type, power, duration, and delay from TTL trigger, as well as logging information **B.** Same as A, but configured to communicate over TCP/IP for multi-language support and broader experimental integration. **C.** Communication schematic for inputs and outputs. Solid lines indicate necessary connections, dashed lines indicate optional connections to and from a data acquisition device (DAQ). Analog outputs (AO, pink) to hardware, and hardware input (red) to PFI0. Zapit PC is directly connected to the DAQ, and can be separate to behavior PC via TCP/IP connection. **D**. The system was programmed to deliver a specific optogenetic stimulus (800 ms with 200 ms rampdown, 40 Hz). A photodiode was positioned at the sample plane to measure photostimulation output (black traces). We also measured the analog input signal sent to the laser (magenta). The inset shows a close up of the photodiode response and analog input signal aligned to the detected onset of the laser input signal (overlay of 50 repetitions). **E**. Low latency and reliable hardware-triggered photostimulation. Optogenetic stimulation (800 ms + 200 ms rampdown, 40 Hz) was triggered using a 100 ms hardware trigger (red trace). Data for each repetition were aligned to the hardware trigger onset. The inset on the right shows a close-up overlay of hardware (red) and photodiode (black) traces aligned to the hardware trigger onset (overlay of 50 repetitions). **F**. Same as E, but showing the photostimulation response when a 1000 ms onset delay was added to the stimulation design. Samples were recorded at 100 kS/s using an oscilloscope (PicoScope)

Zapit supports open-ended and duration-specific stimulation, with controllable stimulus features including stimulus duration, delay, and laser power. Usage of the API is documented on GitHub and examples are provided. Stimulation can be initiated immediately or deferred until a TTL trigger is received, which maximizes timing precision (0.5 ms onset latency and near zero jitter, Figure 6E). Stimuli can also be presented with a fixed delay following the TTL pulse, allowing for precise stimulus alignment with different temporal epochs of a behavioral task (Figure 6F). Zapit has the capacity to write a log file listing each stimulus presentation, so the order of events can be reconstructed *post-hoc*. This is particularly useful in scenarios where the user asked Zapit to produce randomly chosen stimuli. Finally, control trials are possible where the galvos move and the optional masking light is on, but the photostimulation laser is off. Although MATLAB is required to run the Zapit GUI and calibrate the system, experiments can be conducted using any desired programming language. Zapit can be controlled from either the local PC or a remote PC using TCP/IP communication. The Zapit zapit_tcp_bridge package contains Zapit clients for MATLAB, Python, and Bonsai. There is detailed documentation on the message protocol, allowing users to write clients in the language of their choice.

### 2.6 Beam Positioning Accuracy

In trials where ≥2 points are stimulated, the beam must be disabled (blanked) while traveling between locations to prevent off-target stimulation (Figure 7A-B). For simplicity, we chose a fixed motion time of approximately 0.5 ms between any pair of positions. This was long enough to significantly reduce sound generated by the scanners during experiments. The blanking period of the laser was manually tuned to correspond with the motion epoch (Figure 7C) and can be adjusted by the end user for different lasers and scanners. The extent of movement for the XY scanner will change depending on the number of points to be stimulated within a trial and the distance between individual points (Figure 7D).

**Figure 7.**
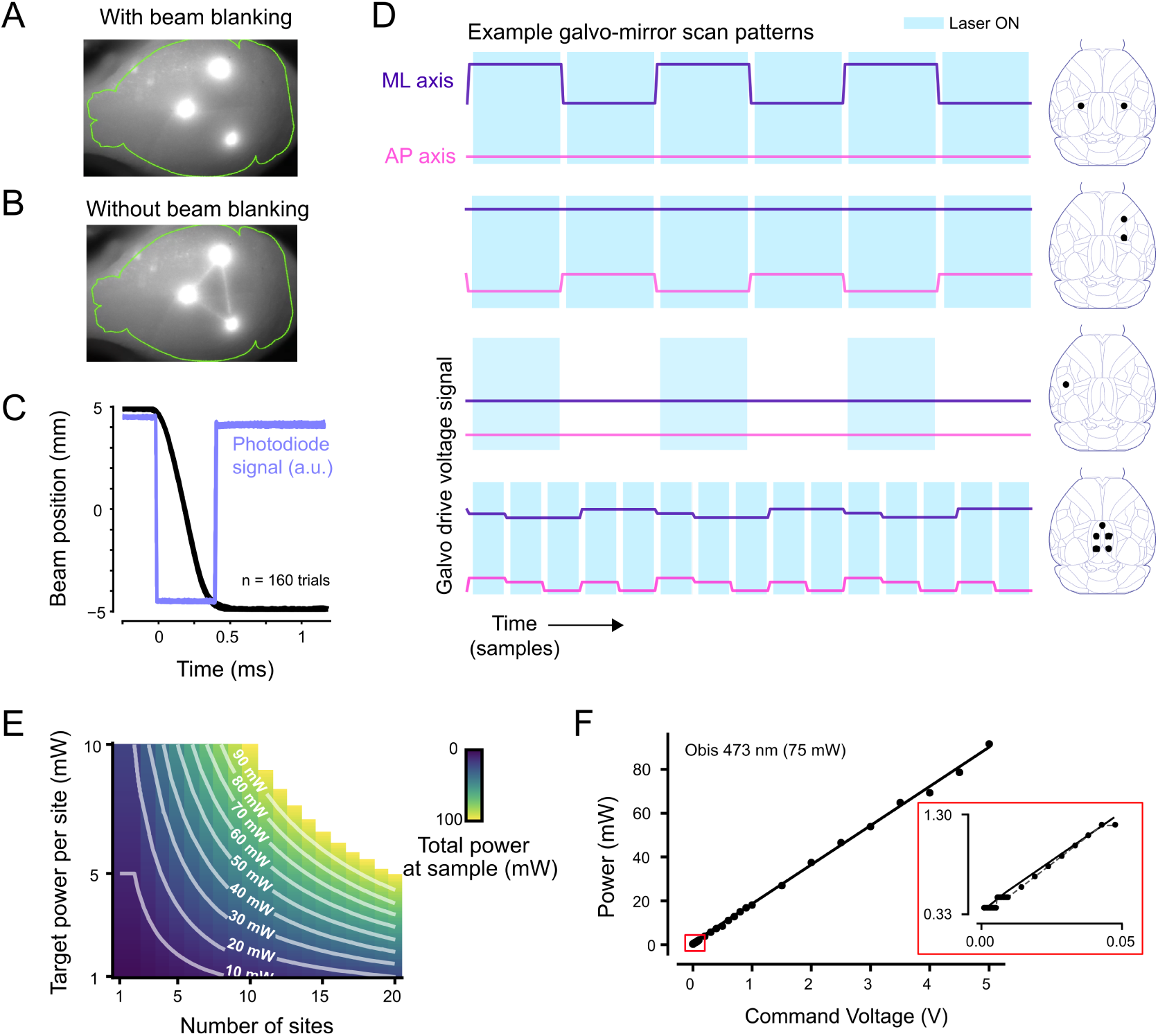
Beam blanking and pointing reproducibility. **A**. Photostimulation of three points with the beam blanked (switched off) as it travels between them. The imaged laser spots are saturated so their size does not reflect stimulation area. **B**. Photostimulation of the same three points but with beam blanking disabled. This leads to visible lines traced over the skull linking the three stimulation points. Blanking avoids this and lowers the risk of off-target stimulation. **C.** Shows the time course and reproducibility of beam pointing and blanking. The beam is cycled back and forth over a 10 mm distance while pointing at a photodiode. The beam is blanked while traveling. We record actual scanner position (black traces) and the photodiode signal (blue traces) displaying negative-going (+5 to −5 mm) transitions. There are 160 overlaid trials. The beam is blanked for about 0.4 ms during the scanner motion epoch. Blanking is consistently perfectly synchronized to scanner motion. The blanking period parameters can easily be tuned for any set of scanners via Zapit’s configuration file. **D.** Example galvanometric mirror scanning patterns for each stimulation paradigm on each axis. Laser onset and offset times are artificially adjusted to emphasize blanking during mirror movement. Blanking periods can be adjusted by the user in system settings. **E.** Theoretical power output by Zapit, calculated by need for target power per site and number of sites, up to 100 mW. **F.** Linear relationship between command voltage and power output for the laser. A linear fit was applied to the data in this case, but Zapit includes functionality for non-linear calibration curves also. Inset: Zoom (red square) to show nonlinearity at ~0 V, with linear model fit from the full data.

The number of sites per trial which can be effectively inactivated will depend on the available laser power. A reference heatmap (Figure 7E) shows an example of the estimated power per site relative to the number of sites.

We chose the Obis laser for its reliability, rise/fall times of µs, and linear power-voltage output. The Obis has sufficient power to maintain an effective average power across many (e.g. 20) sequentially targeted locations. We implemented blanking and power control directly using the Obis laser’s analog control input. This removes the added cost and complexity of an external modulator at the cost of a small non-linearity around 0 mW (Figure 7F). External EOMs and AOMs are supported, via Zapit’s sigmoidal calibration curve option (see Laser Power Calibrations in the GitBook user guide). Even with a sinusoidal stimulation waveform, this discontinuity can create photoelectric artifacts with some electrodes, but does not have a significant physiological impact.

For comparison, we characterized reliability, stability and linear power-voltage output of other lasers (Supplemental Fig. S5), as researchers requiring a smaller number of laser spots per trial could use a cheaper option. When considering the maximal laser power required for an experiment, it is worth keeping in mind losses along the optical path, which can be high in certain circumstances (e.g. some scan mirrors have 20% to 30% losses at 488 nm).

### 2.7 Electrophysiological validation of optogenetic effects

To determine the spatial extent of neural modulation after light propagates through the skull, and relationship to laser power, we performed electrophysiological recordings during photostimulation (Figure 8A-B). We performed these validation experiments in awake VGAT-ChR2-EYFP mice which selectively express the excitatory opsin Channelrhodopsin-2 (ChR2) in inhibitory interneurons. Light stimulation therefore increases local cortical inhibition resulting in suppression of excitatory pyramidal neurons.

**Figure 8.**
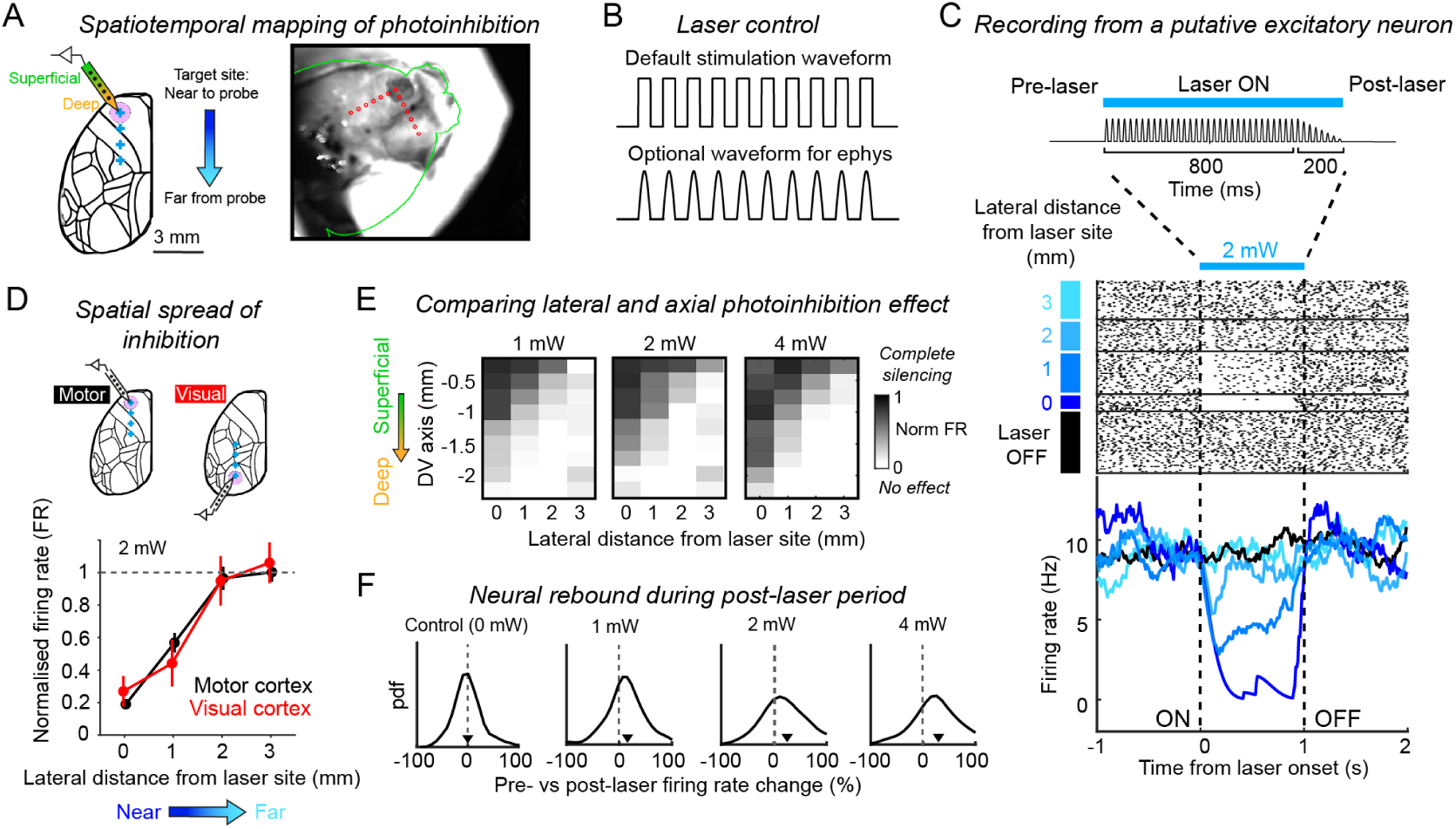
Electrophysiological validation. **A.** Left, neural activity was recorded in awake in VGAT-ChR2-EYFP mice using Cambridge Neurotech silicon probes. Optogenetic effects were characterized as a function of lateral distance (targeting the laser at 0-3 mm away from the recording site). Right, Example image from the Zapit GUI showing a calibration experiment. Red points indicate target sites. A small craniotomy can be seen as the dark patch in the anterior portion of the left hemisphere. **B**. A half-sinusoid stimulation waveform can mitigate opto-electric artefacts during ephys experiments. This waveform option can be implemented by setting the ‘ephysWaveform’ attribute to ‘true’ in the stimulus config file. **C.** Spike raster (top) and peristimulus time histogram (PSTH, bottom) from a putative excitatory neuron in secondary motor cortex (M2) on Zapit and control trials. Zapit trials are sorted according to different lateral distances between the laser target site and the probe location. Photostimulation comprised 800 ms with additional 200 ms linear rampdown. **D.** Suppression of spiking activity as a function of laser beam proximity to the recording site from calibration experiments in M2 (black) and visual cortex (V1, gray). Data show mean and 95% CI. across n = 117 M2 putative excitatory cells and n = 23 V1 putative excitatory cells during 2 mW laser trials. Data are from the same VGAT-ChR2-EYFP mouse. **E.** Suppression of spiking activity across cortical depths and lateral distances for different laser powers. Data show mean across n = 354 putative excitatory cells recorded in M2 (n = 2 VGAT-ChR2-EYFP mice). **F.** Distribution of rebound in firing rates across the M2 excitatory neuron population at different laser powers. We only analysed trials where the laser spot was targeted 0 - 1 mm from the recording site. Rebound was calculated as percentage of firing rate change during the 500 ms following laser offset, relative to pre-laser baseline period. This metric was also calculated for control trials for comparison (black). Arrows indicate the median of each distribution. Data reanalysed from Gauld *et al*. 2025.

To quantify the spatial spread of inhibition, we systematically varied the position of the laser beam across trials at distances ranging from 0 to 3 mm relative to the recording site. On laser-trials, putative excitatory neurons exhibited distance-dependent suppression of spiking that was time-locked to laser presentation (Figure 8C). Suppression was strongest when stimulation was delivered near the recording site and decreased progressively with increasing distance. However, we note that these silencing effects close to the recording site will be exacerbated by the necessity to remove skull to provide electrophysiological access to the brain. With 2 mW time-averaged laser power, we observed no change in firing rate ≥ 2 mm from the recording site. This spatial profile was consistent across cortical areas examined (Figure 8D), and comparable to previous reports of functional resolution using laser-scanning photostimulation (Li *et al*. 2019, Pinto *et al*. 2019). Optogenetic suppression of spiking also decreased as a function of cortical axial depth, with weaker suppression observed ≥ 1 - 1.5 mm from the cortical surface (Figure 8A,E). As expected, both the axial and lateral spread of photoinhibition increased with higher laser power (Figure 8E), with robust silencing effects measured 2 mm below the brain surface using higher power stimulation. Depending on the thickness of cortex at the stereotaxic location of stimulation, this means that deeper cortical and/or subcortical brain areas could be directly impacted with high-power stimulation conditions in transgenic mice with brain-wide opsin expression.

We also quantified the neural ‘rebound’ response following laser offset. Rebound refers to the transient increase in firing rate above baseline levels that can occur following cessation of optogenetic photocurrents. These effects can be mediated via intrinsic biophysical or synaptic properties and/or complex network effects as the circuit recovers from perturbation. Rebound responses are dependent on stimulation parameters, such as duration, laser attenuation duration (ramp-down), and laser power, as well as opsin and transgenic mouse line (Li *et al*. 2019). To assess post-laser changes in firing rate in our calibration experiment, we compared the relative change in firing rate across the pre- and post-laser periods (500 ms; Figure 8C Top). Consistent with previous reports, we observed power-dependent increases in rebound activity, with significant rebound effects at all laser powers above 1 mW (Figure 8F).

Combined, our data suggest limiting stimulation to 1-2 mW (time-averaged power) to minimize spatial spread of inhibition and rebound activity. If possible, users should perform calibration recordings under their own experimental conditions, which could deviate significantly from those shown above.

### 2.8 Behavioral verification of optogenetic effects

The ultimate purpose of the Zapit system is to facilitate *in vivo* optogenetic manipulation experiments that reveal causal relationships between brain activity and behavior. To demonstrate the efficacy of our system in achieving this goal, we performed behavioral validation experiments across multiple research groups and brain regions.

#### Delayed-response somatosensory discrimination

To establish whether Zapit could causally dissociate spatial and temporal neural processes, we trained mice on a delayed-response sensory discrimination task (Figure 9A; Gauld *et al*. 2024). Head-fixed mice received bilateral whisker stimulation (Stimulus epoch; 1 s), and reported the side receiving the stronger stimulus with directional licking following a short delay (Delay epoch; 1 s; Figure 9A-i). In expert mice, we causally tested 17 bilateral sites spanning the dorsal cortex (Figure 9A-ii). On 50% of trials, a single bilateral site was stimulated during either the Stimulus or Delay epoch. Mice performed with high accuracy on control trials without laser stimulation (P(Correct) = 0.89 + 0.08; mean ± std across 76 sessions from 4 mice). However, consistent with previous work (Guo, Li, *et al*. 2014), we found epoch- and site-specific photoinhibition effects on task performance (Figure 9A-iii). During the Stimulus epoch, only inhibition of barrel cortex and anterolateral motor cortex (ALM) significantly impaired correct choices. Barrel cortex is the area of primary somatosensory cortex (S1) that receives input from the whiskers, and ALM is a premotor cortical area involved in planning tongue movements, so inactivating these regions during the sensory processing epoch is expected to impair the animal’s ability to perform the task. During the delay epoch, we found particularly robust effects of ALM inactivation on choice accuracy and reaction time, consistent with this area being critical for motor planning and short-term memory maintenance (Guo, Li, *et al*. 2014, Li *et al*. 2016). We then performed unilateral inactivation experiments targeting ALM and observed robust ipsilateral choice biases, consistent with contralateral hemispheric motor-processing (Figure 9A-iv). As expected from our electrophysiological quantification, behavioral effects were power-dependent, saturating at ~2 mW (Figure 9A-v). Zapit was therefore able to confirm epoch-specific causal links between different cortical areas and distinct behavioral processes.

**Figure 9.**
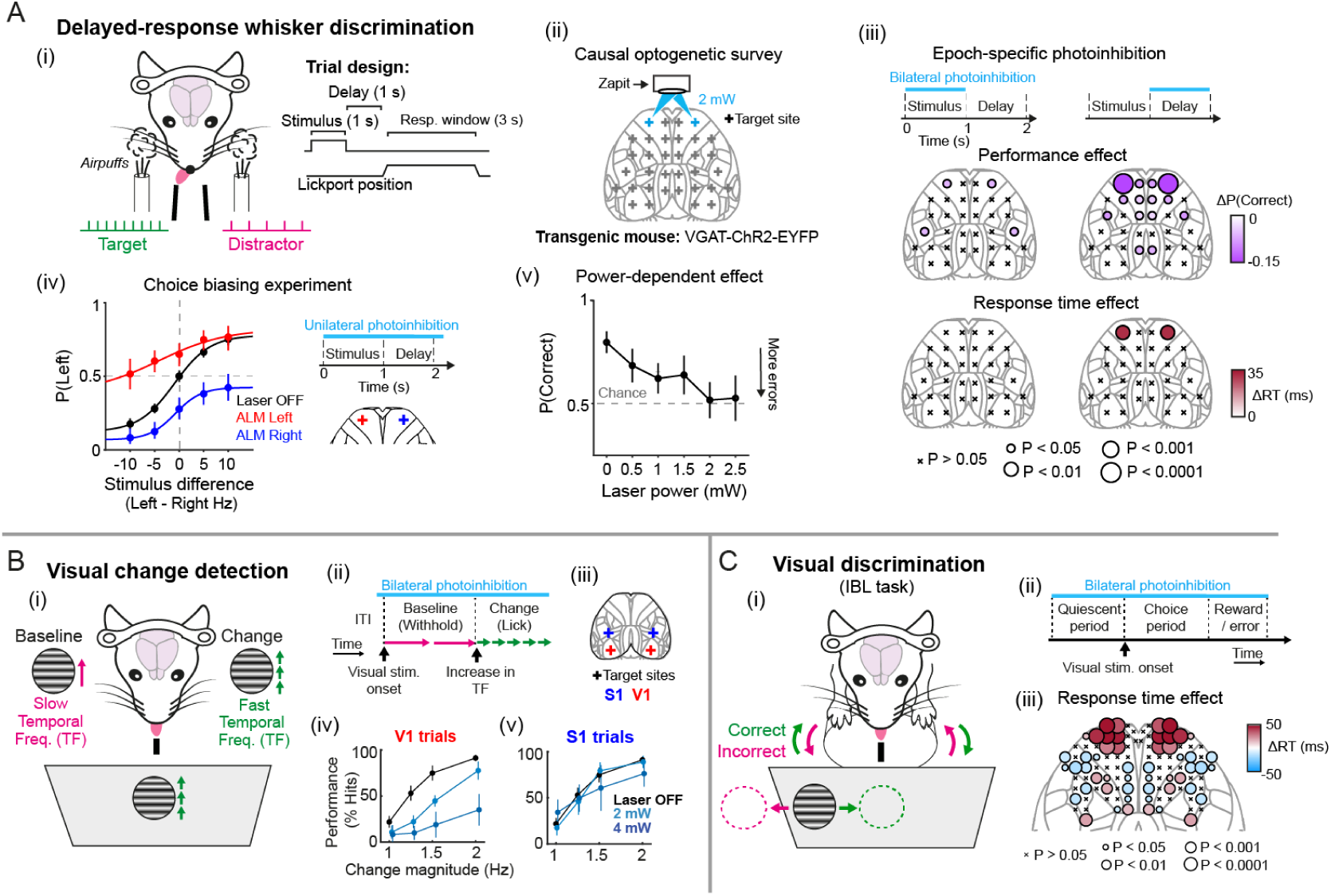
Behavioral validation experiments. **A**. Whisker discrimination **(i)** Overview of task and trial design. Mice discriminated bilateral whisker input and reported decisions with directional licking following a delay. A motorized lickport moves in/out on each trial to cue the response window. **(ii)** Summary of bilateral Zapit target sites. **(iii)** Results from photoinhibition experiments in expert mice. The maps show the mean effect (color) and statistical significance (circle size) of site-specific photoinhibition on choice accuracy (top) and reaction time (bottom), during the stimulus (left) and delay (right) epochs. ‘X’ indicates non-significant effects (P > 0.05). N = 76 sessions, 4 VGAT-ChR2-EYFP mice. **(iv)** Results from unilateral anterolateral motor cortex photoinhibition experiments. Unilateral inactivation resulted in ipsilateral choice biases, as seen by vertical shifts of the psychometric curve. Psychometric curves are shown for control trials (black) and left (red) and right (blue) ALM photoinhibition trials. Data points show the mean and error bars show 95% CI across sessions. N = 11 sessions, 3 mice. **(v)** Dependence of behavioral impairment on laser power. **B**. Visual change detection. **(i)** Overview of task design. Mice viewed a drifting visual stimulus and reported sustained increases in temporal frequency with licking. **(ii)** Overview of trial design and optogenetic stimulus timing. **(iii)** Summary of Zapit target sites. **(iv)** Effect of V1 photoinhibition on task performance. Data show mean and 95% CI across mice (N = 5 mice). Control trials (black), 2 mW (light blue) and 4 mW (dark blue) photoinhibition trials. **(v)** Same as (iv) but for S1 photoinhibition trials. **C**. Visual discrimination. **(i)** Overview of task design. Mice discriminated the spatial location of a visual stimulus and turned a wheel to translate the stimulus to the center of the screen. **(ii)** Overview of trial design and optogenetic stimulus timing. **(iii)** Results from a photoinhibition mapping experiment comprising a grid of 52 bilateral stimulation sites positioned at 0.5 mm intervals. The map shows the mean reaction time on Zapit trials (color), and statistical significance as tested against control trials (circle size). ‘X’ indicates non-significant effects (P > 0.05).

#### Visual change detection

To demonstrate the versatility of Zapit across task designs, we performed a separate set of experiments in mice trained to report changes in the temporal frequency of a drifting visual stimulus (Figure 9B-i-iii; Orsolic *et al*. 2021). On control trials without laser stimulation, mice showed near 100% performance when the stimulus change was easy to detect. As expected for a visually-guided task, bilateral inactivation of primary visual cortex (V1) produced clear deficits in performance, with higher laser power resulting in stronger impairment (Figure 9B-iv). In contrast, inactivation of S1 did not significantly impact behavior (Figure 9B-v, consistent with the task having no somatosensory component). However, we did observe a small, but non-significant, decrease in psychometric performance during high power (4 mW) S1 photoinhibition (Figure 9B-v dark blue), which could be a result of unintentional off-target light ‘spillover’ impacting nearby higher-order visual areas. We therefore show that Zapit can also be used to generate robust and selective impairments in perceptual processing during visually-guided behavior.

#### Visual discrimination

To explore the effective spatial resolution of this inactivation approach with Zapit, we implemented a fine-scale causal mapping experiment in mice tasked to report the location of visual stimuli by turning a wheel (Figure 9C-i,ii; International Brain Laboratory *et al*. 2021; International Brain Laboratory *et al*. 2025). The stimulus-set comprised 52 bilateral sites positioned at 0.5 mm intervals covering a large network of frontal, motor, and somatosensory cortical areas. Photoinhibition of frontal motor cortex sites produced robust increases in reaction time (Figure 9C-iii left). In contrast, stimulation at other motor and somatosensory sites produced modest but significant decreases in reaction time. Our findings therefore demonstrate site-specific and opposing effects on the control of motor behavior across distributed cortical networks.

Together, these behavioral experiments demonstrate that Zapit enables precise and inter-pretable causal mapping of cortical contributions to behavior.

## 3 Discussion

Zapit is a complete and accessible solution for laser-scanning optogenetics. The components are commercially available, and the modular system is uniquely flexible to accommodate a diversity of experimental needs. The Github repository contains complete build instructions, parts lists, and visual aids: no prior expertise is required for implementation. The software is fully documented and will be continually maintained and updated by developers and experimentalists. Known software and hardware issues are actively raised and resolved through Zapit’s Github issues page.

Zapit comprises several technical developments that make it more versatile and user-friendly than any previously published system. These include detailed configurations for multiple lasers and opsins, user-friendly GUIs for automated stereotaxic calibration and stimulus design, and a cross-platform API for integration into diverse behavioral and experimental protocols across laboratories. Additionally, the modular design facilitates future developments, such as multiplexing two lasers in one setup.

Zapit produced robust behavioral effects across a variety of experimental protocols and brain regions with VGAT-ChR2 mice. We validated the efficacy of inactivation in this mouse strain with simultaneous electrophysiology recordings and demonstrate that the effective radius of inactivation depends on the laser power used, but was ~1 mm at powers for which we observed significant behavioral perturbations.

There are other effective approaches for rapid programmatic photostimulation in head-fixed behavioral tasks: those based on digital micro-mirror devices (DMDs – Chong *et al*. 2020) or spatial lights modulators (SLMs – Russell *et al*. 2022; Gauld *et al*. 2024). DMD-based solutions, such as the Mightex Polygon, provide a faster pattern-switching time but due to light loss, these systems require *~100 times* more laser power than Zapit to cover the mouse dorsal cortex. SLM-based solutions are more light efficient, but devices with high resolution are very expensive and are slower than DMDs. Therefore, Zapit’s scanner-based approach provides the most cost-effective and compact solution for the majority of optogenetics experiments involving the dorsal cortex.

### 3.1 Practical considerations when using Zapit

#### Surgical procedures

To maximize targeting flexibility and accuracy, we recommend a head-fixation device that provides an unobstructed view of the entire dorsal surface of the skull, and marking structural reference points during surgery. We further recommend a thin layer of transparent glue (Zatka-Haas *et al*. 2021; Coen *et al*. 2023) or ‘bone’ cement (200-300 *µm* thick) to protect the skull, as thicker layers of cement (0.5-1 mm) may reduce stimulation efficacy. For general details of *in vivo* surgical and behavioral procedures in head-fixed mice we direct the reader to existing resources (e.g. Guo, Li, *et al*. 2014; Guo, Hires, *et al*. 2014).

#### Laser and power selection

Zapit is compatible with any laser source that supports fast analog power modulation. As a guideline, for VGAT-ChR2 mice we recommend 1–2 mW time-averaged power at the skull surface. With this line and these powers we consistently observe strong behavioral effects, with spatiotemporal resolution comparable to that reported in other studies. Light loss through the skull and bone cement is around 50% (Guo, Li, *et al*. 2014). Heating at these powers is minimal, reducing the risk of heat-induced changes in neural activity (Stujenske, Spellman, and Gordon 2015). Nevertheless, *post hoc* histology is strongly recommended to assess tissue integrity and confirm opsin expression. Regular power measurements at the sample are essential for stability and reproducibility.

#### Targeting precision

With transgenic lines expressing opsin brain-wide (including VGAT-ChR2 mice), stimulation with high laser powers (e.g. 4 mW or above at the sample) may cause off-target behavioral effects due to light-scatter laterally and axially through cortex (Li *et al*. 2019; Babl, Rummell, and Sigurdsson 2019). Ideally, laser parameters including power, duration and ramp-down should be informed by electrophysiological recordings and/or modeling of light propagation through the brain (Yona *et al*. 2016). If limiting the impact of light dispersion is critical, opsins can be expressed via targeted viral injections. Under these conditions, expression can be restricted to highly localized circuits, genetically and anatomically defined subpopulations and sub-cellular compartments, and verified using *post-hoc* histology.

#### Sensory confounds

Fast movements of galvo-mirrors generate auditory cues that can influence behavior, and we therefore recommend moving the galvo-mirrors on all trials, including those where the laser power is 0 mW. Zapit behaves this way by default. The galvo drive waveforms are shaped to reduce noise emission, and the scanner unit can also be enclosed in sound-attenuating housing to further mitigate these effects. To accurately interpret optogenetic behavioral effects, it is also critical to mask the mouse’s ability to see the laser, whether through internal and/or external retinal stimulation (Danskin *et al*. 2015; Weiler *et al*. 2026). We recommend masking laser-emitted light with color-matched LEDs placed above the mouse on every trial, matched to the stimulation profile of the laser (e.g., 40 Hz), and using a well-lit setup. Extra caution should be taken when using longer wavelengths which easily propagate through brain tissue to the retina, and can be more difficult to mask (Danskin *et al*. 2015; Weiler *et al*. 2026). Sensory confounds can also be addressed through repeated experiments and analyses in control animals that do not express the opsin (Gauld *et al*. 2025).

#### Target region selection and interpretation

A major advantage of Zapit is the capacity to rapidly survey causal effects across the entire dorsal cortex. However, skull curvature and thickness should be considered, particularly in more lateral cortical regions. Differences in opsin expression due to tropisms in viral vectors or transgenic lines can also influence results. Users may need to perform further experiments to determine whether observed effects are a direct result of regional inactivation, or derived from downstream brain regions (Wolff and Ölveczky 2018; Allen 2017).

### 3.2 Good practice: measures to report when using Zapit

We encourage Zapit users to report a minimal set of standardized parameters to aid replication, comparability, and consistency across experiments and laboratories:

- Average light power per site
- Stimulation frequency and profile (e.g. 40 Hz sinusoid), and duty cycle
- Laser stimulation and ramp-down duration
- Specifications and duty cycle of any masking light
- Precise stimulation coordinates

### 3.3 In Closing

We hope Zapit is a tool that will grow and find a broad user base throughout the scientific community. Potential new users are welcome to contact us for assistance in building or setting up the system.

## 4 Methods

### 4.1 Code

Zapit and its associated software tools were developed in the open, version controlled with Git, and hosted on GitHub. Most development was done using Sublime Text. Anthropic Claude (Fable 5 and Opus 4.7) was used to generate code reviews and identify bugs in Zapit’s 1.0.4 release.

### 4.2 Recording and data acquisition for hardware characterization

Laser power output for all lasers was recorded using a Thorlabs Power Meter PM100D, and acquired with the associated Thorlabs Optical Power Meter Utility software. Samples for photostimulation reliability with the Obis 473 nm laser (Figure 6) were recorded at 100 kS/s using an oscilloscope (PicoScope). Samples for photostimulation reliability for all other lasers were recorded at 100 kS/s using a NI-DAQ USB-6453 and acquired with LabVIEW software. In both cases, laser output was recorded using a photodiode from a programmed 800 ms stimulation period, following a 200 ms rampdown. For users without a LabVIEW license who wish to acquire similar data we provide example code in Zapit (derived from our MATLAB DAQmx examples package) showing how to do so using NI DAQmx via MATLAB. Python NI DAQmx examples are also available.

### 4.3 Zapit configuration for experiments

In all experiments presented here we used a 473 nm, 75 mW Obis laser with a Thorlabs MD498 dichroic filter and an MF525-39 emission filter. The system was housed in a custom enclosure. ScannerMax Saturn 5 mirrors (available from Edmund Optics) were used to direct the beam. However, we recommend the ThorLabs version presented in this paper for most users, as it is more cost effective and easier to build. Other scanners could also be substituted, as can other lasers, lens combinations, and laser wavelengths etc. Zapit was controlled and data were acquired with an NI-DAQ USB-6363.

### 4.4 Mice

All experiments were performed under the UK Animals (Scientific Procedures) Act of 1986 (PPL: PD867676F) following local ethical approval by the Sainsbury Wellcome Centre Animal Welfare Ethical Review Body. We used VGAT-ChR2-YFP mice (Jackson Laboratories, USA) for all experiments.

### 4.5 Surgical procedures

Implantation of a clear skull cap (i.e., thin layer of transparent bone cement (C & B Superbond)) covering the dorsal surface of the brain together with a head plate placed over the cerebellum was carried out under 1.5% isoflurane in O2, together with pre and post surgical administration of Meloxicam (5 mg/kg). To ensure the skull is level under the objective, the skull should be aligned by checking that bregma and lambda depths are even (i.e., <0.05-0.1 mm difference), as well as ± 2mm lateral from midline is even (i.e., <0.05-0.1 mm difference). To allow calibration of the sample relative to stereotaxic space during Zapit experiments, two small dots should be made over bregma and +3 mm AP from bregma.

For electrophysiological validation of Zapit photoinhibition, craniotomies (~ 1.5 mm diameter) were made over secondary motor cortex (AP + 2.5 mm, ML ± 1.5 mm, from bregma) and visual cortex (AP −2.9 mm, ML 1.4 mm, from bregma). An additional cranial opening (~ 0.5 mm diameter) was drilled above the cerebellum to secure the ground wire to the skull. Then the craniotomy was covered with Dura-Gel (Cambridge NeuroTech), and a plastic protective cap was placed over the headplate to shield the exposed areas and maintain sterility between recording sessions.

### 4.6 Electrophysiological recordings

For electrophysiological validation experiments, neural activity was recorded using a 64-channel silicon probe (ASSY-236-H9, Cambridge NeuroTech) to minimize photoelectric artifacts. Signals were digitized at 30 kHz, amplified, and band-pass filtered (0.6–7500 Hz) using a 64-channel Intan Technologies headstage. Data were acquired, visualized, and stored using the Open Ephys acquisition board and GUI. During the recording, probes were lowered at 7 *µ*m/s using programmed automatic micromanipulators (Sensapex), followed by a 10 minute settling period to minimize neural signal drift during recording. At the end of each trial, a serial TTL message encoding the current trial number was sent from behavioural control hardware to the acquisition system to synchronize the neural signal with the Zapit trigger signals.

Offline spike sorting was performed with Kilosort v4 (Pachitariu *et al*. 2024), followed by manual curation in Phy 2.0. Units were classified as putative neurons only if their spike waveforms exhibited a clear spatial decay across adjacent probe channels. Both single and multi units that showed low percentage of missing spikes, low refractory period violations, and minimal drift across time were included in the analysis. Units exhibiting substantial drift that led to a clear loss of spikes over time were excluded from further analysis. Additional details can be found in Gauld *et al*. 2025.

### 4.7 Delayed-response somatosensory discrimination task

The task design and experimental set-up were based on previous work (Gauld *et al*. 2024; Gauld *et al*. 2025). Briefly, mice were head-fixed in a perspex tube and air-puff stimuli (0 - 10 Hz; 20 ms pulse duration, 1 s sequence duration) were delivered to the left/right whiskers. Following a short delay period (1 s), mice reported their choice with directional licking to obtain water rewards (3 *µl*). The target response side was cued by the whisker side receiving the faster air-puff sequence, and randomized on trials with matched bilateral stimulation.

Following task learning, we surveyed 17 bilateral regions spanning the dorsal cortical surface (stereotaxic target coordinates in mm relative to bregma): (2.5 A ± 0.5 L), (2.5 A ± 1.75 L), (1.25 A ± 0.5 L), (1.25 A ± 1.75 L), (1.25 A ± 3 L), (0 A ± 0.5 L), (0 A ± 1.75 L), (0 A ± 3 L), (−1.45 A ± 0.5 L), (−1.25 A ± 1.75 L), (−1.25 A ± 3.5 L), (−2.7 A ± 0.5 L), (−2.5 A ± 1.75 L), (−2.5 A ± 3 L), (−2.5 A ± 4.25 L), (−3.75 A ± 1.75 L), and (−3.75 A ± 3 L). Laser stimulation (time-averaged power per site: 2 mW (at the sample); duration: 800 ms + 200 ms linear rampdown) was delivered on 50% of trials, with target-site and epoch (Stimulus vs Delay) fully randomised. During the optogenetic mapping experiment, mice received unilateral whisker stimulation trials (10 Hz).

To examine putative epoch- and site-specific effects of laser stimulation on task performance, we compared Zapit trials for each bilateral target site with session-matched control trials using generalized linear mixed models (GLMMs). Specifically, we compared the predictive performance of the following two models shown below in Wilkinson-Rogers notation:

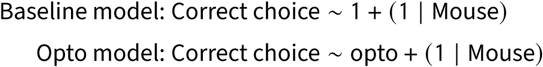

We performed 20-fold cross-validation to evaluate model predictive performance on out-of-sample test data. Mean log-likelihoods across matched folds were compared using a paired *t*-test, yielding a *P*-value for each cortical target site and each epoch. For analyses of reaction times (RTs), we compared Zapit and control trial RTs with a Wilcoxon rank-sum test. For performance and RT analyses, *P*-values were corrected for multiple comparisons across cortical locations using the Bonferroni–Holm method (*α* = 0.05, 17 cortical locations).

For choice-biasing experiments, we selectively targeted anterolateral motor cortex (“ALM”: 2.5 A± 1.75 L) for unilateral photoinhibition. Laser stimulation (time-averaged power per site: 2 mW (at the sample); duration: 2000 ms + 200 ms linear rampdown) was delivered on 33% of trials and randomized across hemispheres. During choice-biasing experiments, mice received nine sensory trial-types corresponding to a 3 × 3 combinatorial matrix of 0, 5, and 10 Hz left and right stimuli. Trials were delivered in randomized order.

### 4.8 Visual change detection task for proof of principle experiments using Zapit

The design of the behavioral task and stages of behavioral training were as previously described in Orsolic *et al*. 2021.

Mice were head-fixed and placed on a polystyrene wheel. Two monitors (21.5”, 1920 x 1080, 60 Hz) were placed on each side of the mouse at approximately 20 cm from the mouse head. The monitors were gamma corrected to 40 cd/m^2^ of maximum luminance using custom MATLAB scripts utilizing PsychToolbox-3. The stimulus presentation was controlled by custom written software in MATLAB utilizing PsychToolbox-3. The visual stimulus was a sinusoidal grating with the spatial frequency of 0.04 cycles per degree resulting in 3 grating periods shown on a screen. Each trial began with a presentation of a gray texture covering both screens. After a randomized delay (at least 3 s plus a random sample from an exponential distribution with a mean of 0.5 s), the baseline stimulus appeared. The temporal frequency (TF) of the grating was drawn every 50 ms (3 monitor frames) from a log-normal distribution, such that log_2_-transformed TF had the mean of 0 and standard deviation of 0.25 octaves and a geometric mean of 1 Hz. The direction of drift was randomized trial to trial between upward or downward drift. The sustained increase in TF, referred to in the text as change period, occurred after a randomized delay (3-15.5 s) from the start of baseline period and lasted for 2.15 s. Random 15% of trials were assigned as no-change trials and did not have a change period.

Mice were trained to report sustained increases in temporal frequency by licking the spout to trigger reward delivery (drop of soy milk). Licks that occurred outside of the change period are referred in the text as early licks. If mice moved on the running wheel (movement exceeding 2.5 mm in a 50 ms window) in either direction, the trial was aborted. If mice did not lick within 2.15 s from the change onset, the trial was considered a miss trial.

Optogenetic silencing commenced 0.25 s before visual stimulus onset using 40-Hz square-wave stimulation (50% duty cycle) delivered bilaterally throughout each trial. Following trial completion, laser power ramped down linearly over 0.25 s to minimise rebound excitation. To ensure mice could not distinguish laser from control trials based on visual cues, we combined two masking strategies. First, we illuminated the behavioural chamber with white LED strips throughout all sessions. Second, we delivered blue LED light from above the animal on every trial, matching the exact temporal profile of the optogenetic laser stimulation. This dual masking approach prevented mice from using visual detection of laser light to alter their behaviour.

We silenced primary somatosensory cortex (S1: ML=±3.2, AP=-0.8 mm) and primary visual cortex (V1: ML=±2.7, AP=-3.8 mm). Each area was silenced bilaterally using either 2 or 4 mW average power with each power level applied on 3.6% of trials per site. We also silenced five additional sites in these sessions not reported in this paper. 50% of trials served as no silencing controls where the laser was either turned off or targeted away from the skull.

Psychometric curves were calculated for each mouse by counting the number of hits relative to all trials where mice were presented with a change stimulus (i.e., did not early lick nor abort), across all sessions. Mean hit rates (performance) were calculated for each mouse (n = 5) per change size for either intact cortex (No area silenced) or for trials silencing each area with each light power. To ensure fair aggregation of trials across sessions, we only included sessions with performance above 80% hits on largest change (2 Hz) in cortex intact trials. We plotted the mean and 95% confidence intervals (i.e. standard error of the mean (s.e.m.) * 1.96) of hit rates across mice for each change size and each condition. We quantified differences between area silencing trials and intact cortex trials by performing paired t-tests (across mice) on the combined difference in performance for change sizes 1.25 and 1.5 Hz for intact cortex vs a specific area silenced (excluding 2 Hz to avoid potential statistical selection bias).

### 4.9 Visual discrimination task

Surgical procedures and behavioral training followed established protocols (International Brain Laboratory *et al*. 2021; International Brain Laboratory *et al*. 2025) with minor modifications. Mice were water-restricted and trained on the 2-AFC task using standardized training rigs. Mice were transitioned to the laser-scanning optogenetics (Zapit) rig upon reaching the “training 2b” performance criterion. The behavioral apparatus was adapted from the standard IBL electrophysiology rig design (International Brain Laboratory *et al*. 2021).

A grid of 52 bilateral stimulation sites was positioned at 0.5 mm intervals to cover secondary motor cortex, primary motor cortex, the upper limb region of somatosensory cortex as well as parts of anterior cingulate and prelimbic cortices.

Laser stimulation was controlled via TTL signals generated by modified IBL training software. On stimulation trials, TTL signals were sent at quiescent period onset and trial end (reward, error, or timeout). On each trial, there was a 50–70% probability that one bilateral site pair would be randomly selected for stimulation. Generally, the system was run with 2 mW laser power, with some exceptions for sessions in which 3 mW was used. To control for visual artifacts, a 473 nm LED masking light was positioned near the laser-scanning source and was activated on every trial coincident with the laser stimulation window, regardless of whether actual stimulation occurred.

## 5 Author Contributions

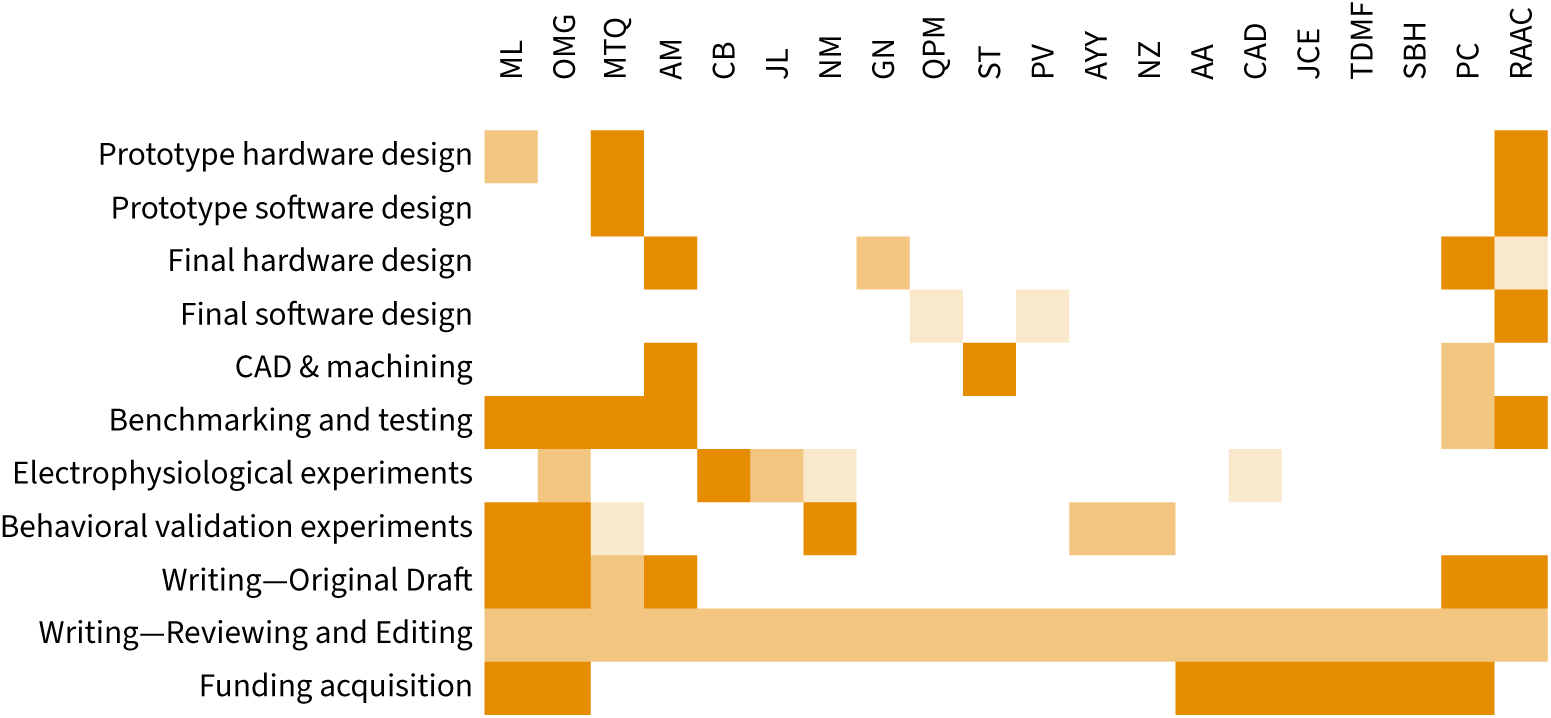

## Acknowledgments

We thank Andy Peters for providing us with a MATLAB script for displaying a top-down view of the Allen Atlas in stereotaxic coordinates, and inspiring us with the Neuropixels Trajectory Explorer (10.5281/zenodo.7043459). We thank Charu Reddy and Matteo Carandini for sharing resources to facilitate development, and Michael Krumin and Simon Weiler for valuable conversations. Dale Elgar from COSYS Ltd. did most of the custom enclosure design for additional acoustic attenuation. Graeme McPhillips at the SWC Electronics Core Facility provided advice on construction of the electronics enclosure. We thank Morio Hamada and Ivan Voitov for providing feedback on the manuscript.

This work was supported by Wellcome awards to T.D.M.F. (217211/Z/19/Z), M.L. (224121/Z/21/Z), O.M.G. (309104/Z/24/Z) and P.C. (225992/Z/22/Z). C.A.D. was supported by UKRI grant EP/Y008804/1. HF, NM, and AY were supported by The Wellcome International Brain Laboratory award (216324/Z/19/Z) and the Simons Culmination award (SFI-AN-NC-GB-IBL-00002672-07). The Advanced Microscopy Facility and individual labs were also supported by the Sainsbury Wellcome Centre’s core provided by Wellcome (219627/Z/19/Z) and the Gatsby Charitable Foundation (GAT3755).

## Data Availability Statement

This project is developed entirely in the open: all code is available at github.com/Zapit-Optostim. Code and processed data required to re-create our figures is shared at github.com/Zapit-Optostim/overleaf-manuscript. Raw data files underlying these can be provided on request.

## 6 Supplementary Figures

**Figure S1.**
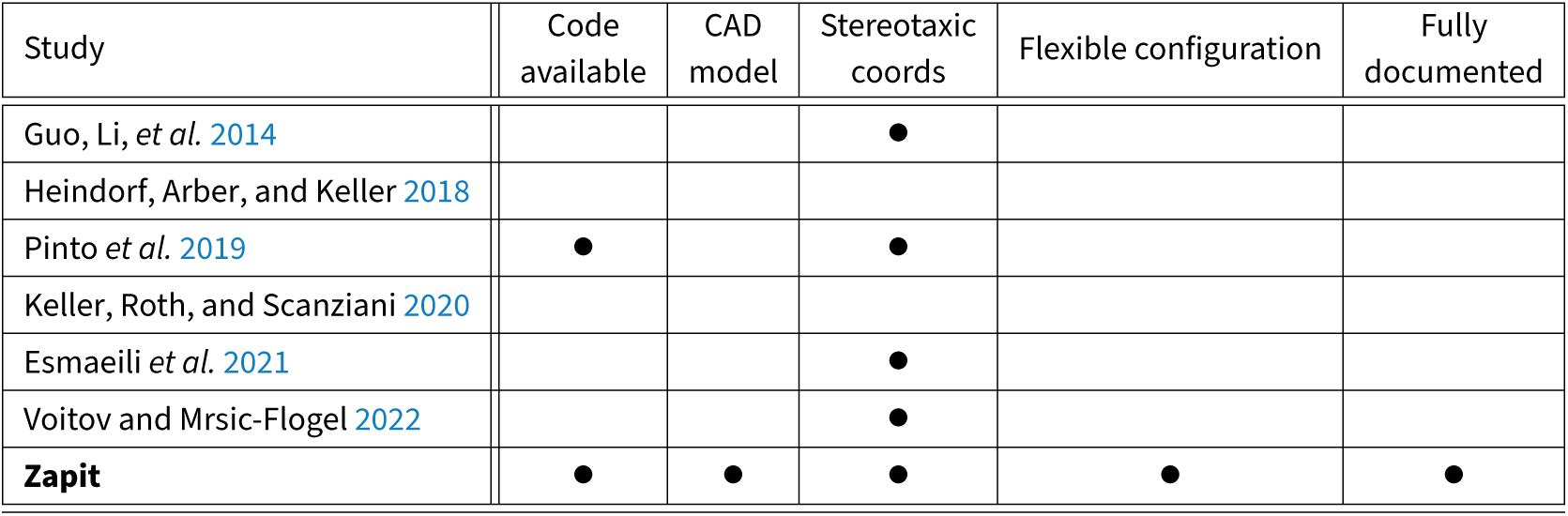
Comparison of Zapit to related systems. ‘*Codeavailable*’ indicates whether readers can easily download code capable of running the author’s random-access optostimulation paradigm. Only Pinto *et al*. 2019 provide a clear standalone codebase for running their optogenetic system. Heindorf, et al. have shared their code embedded within their LabVIEW-based multiphoton scanning software, Iris, but this is not documented. Also, while all previous systems are of high quality, they are very specific to the published experimental protocol and so do not have a ‘*Flexible configuration*’. No published study has an associated *CAD model* or build instructions. Not all studies specify coordinates to be stimulated in stereotaxic space. As far as we know, only Zapit has a GUI for building stimulus sets in sterotaxic space.

**Figure S2.**
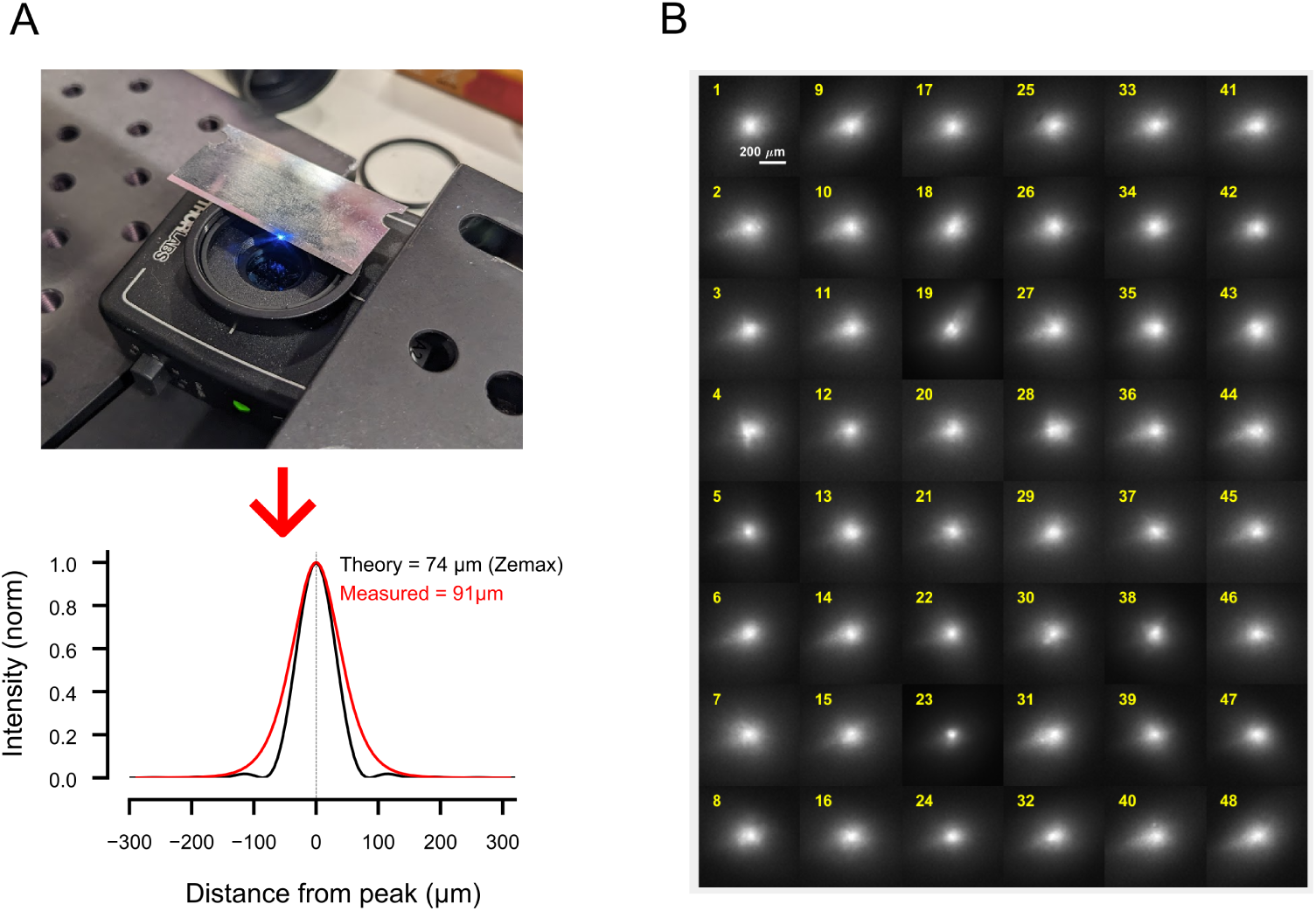
Resolution of the scanning system. **A**. Theoretical and measured X/Y PSF based on a f=150 mm objective lens, a 0.8 mm beam and a Coherent Obis 473 nm 75 mW laser. The empirical data were obtained by translating the focused laser spot over the edge of a razor blade and measuring light intensity with a photodiode placed under the blade. This procedure yields an intensity curve resembling a cumulative Gaussian, which can be fitted as such and converted to a probability density function. While we measure a beam size of under 100 µm FWHM, the true size of the spot on the brain surface is likely to be much larger. The beam will scatter as it goes through the cleared skull and then will scatter further as it enters the brain. **B**. Beam spot size across the field of view. Focusing the beam on a piece of paper elicits fluorescence which can be imaged with the camera. We acquired many such images while moving the beam over a grid of positions spanning an area roughly the size of a mouse brain (14.0 mm by 10.5 mm, leading to a 1.75 mm separation between points). The image is a composite showing zoomed-in details of the laser spot over all 48 beam locations. The image tiles are discontinuous so the scale bar refers to the beam shape and does not provide information about the separation between adjacent tiles. The size and shape of the beam is very similar across all positions, showing there is no change in the resolution of optical stimulation over the field of view.

**Figure S3.**
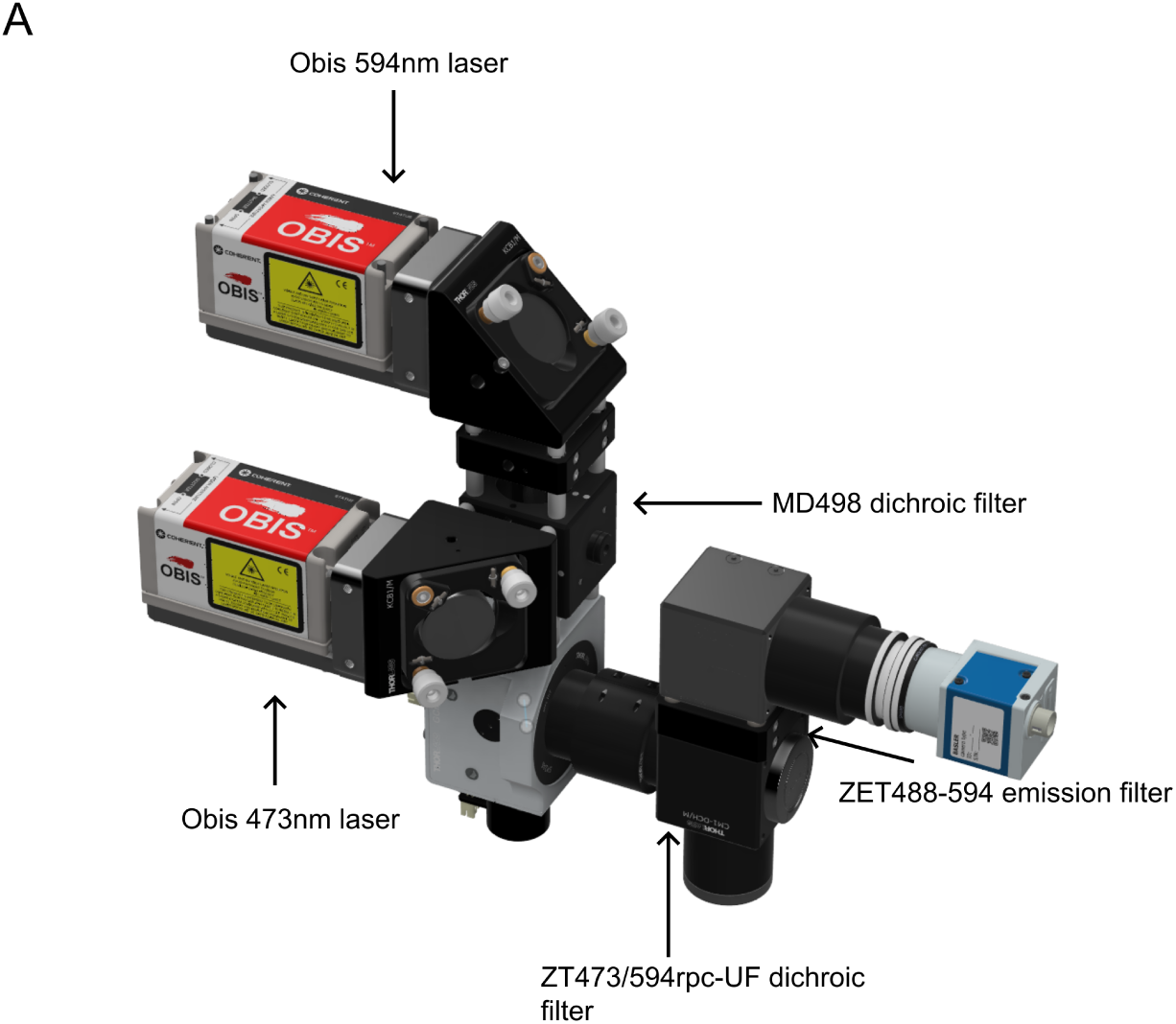
Dual-laser configuration for multiple opsin stimulation and real-time imaging. **A.** The Zapit system can be assembled to mount two lasers, both of which can be controlled within one experiment. An additional dichroic (Thorlabs) is needed to merge the two beams before the scanners. Resulting fluorescence passes through the chosen emission filter (Chroma). Ongoing changes to Zapit will allow for dual-laser control in the future.

**Figure S4.**
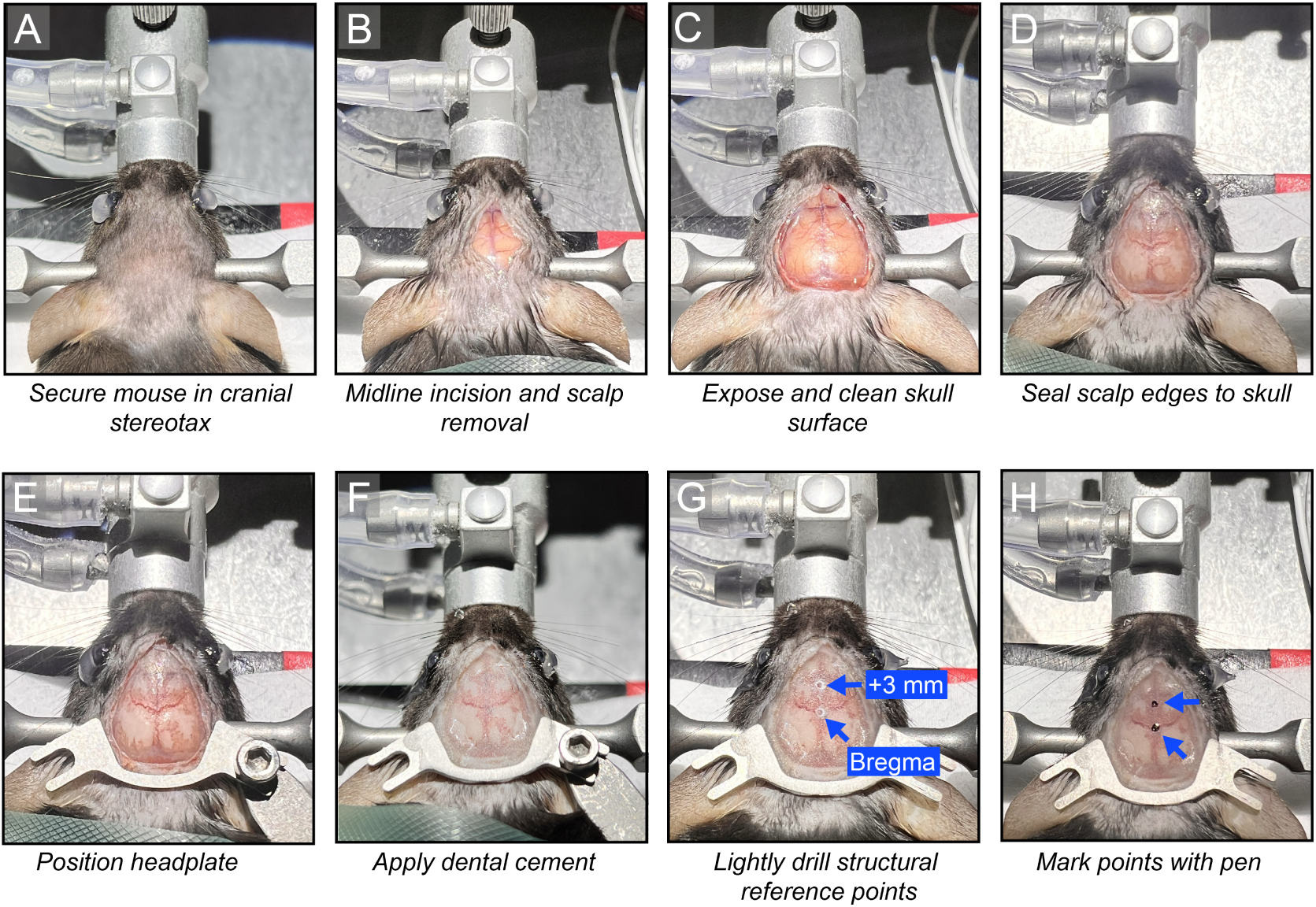
In vivo surgical procedures for Zapit-based experiments. All steps should be performed under sterile conditions with appropriate pre- and post-operative care procedures in place. **A**. An anesthetized mouse is installed in a cranial stereotax via bite and ear bars. Inhalation anesthesia is provided via a nose cone. Prior to this step, the scalp is shaved and cleaned to prevent fur contaminating the surgery area during incision. **B**. An incision is made along the midline of the scalp and a small section of scalp is removed bilaterally using surgical scissors. **C**. The edges of the scalp can be gently moved to the sides using sterile swabs to expose the dorsal surface of the skull. Sterile saline and swabs can be used to clean the skull and any areas of bleeding. A bone-scraping tool can be used to remove periosteum/membrane on the surface of the skull. This is critical to ensure good adhesion of bone cement to secure the headplate. **D**. The edges of the scalp are gently fixed to outer edges of the skull using tissue adhesive, ensuring a large area of skull remains exposed. **E**. A custom metal headplate is then positioned at the back of the animal’s skull. This headplate is designed to provide unobstructed optical access to dorsal cortex. **F**. The headplate is secured with transparent bone cement. A thin layer of cement is applied across the dorsal skull surface to seal the area and prevent infection. **G**. Two structural references points are marked in the cement once cured using a small drill. The points correspond to bregma and a position 3 mm anterior from bregma. **H**. Visibility of the reference points is increased with marker pen to ensure clear identification when viewing the sample through the Zapit system camera. This is critical for accurate sample calibration and alignment.

**Figure S5.**
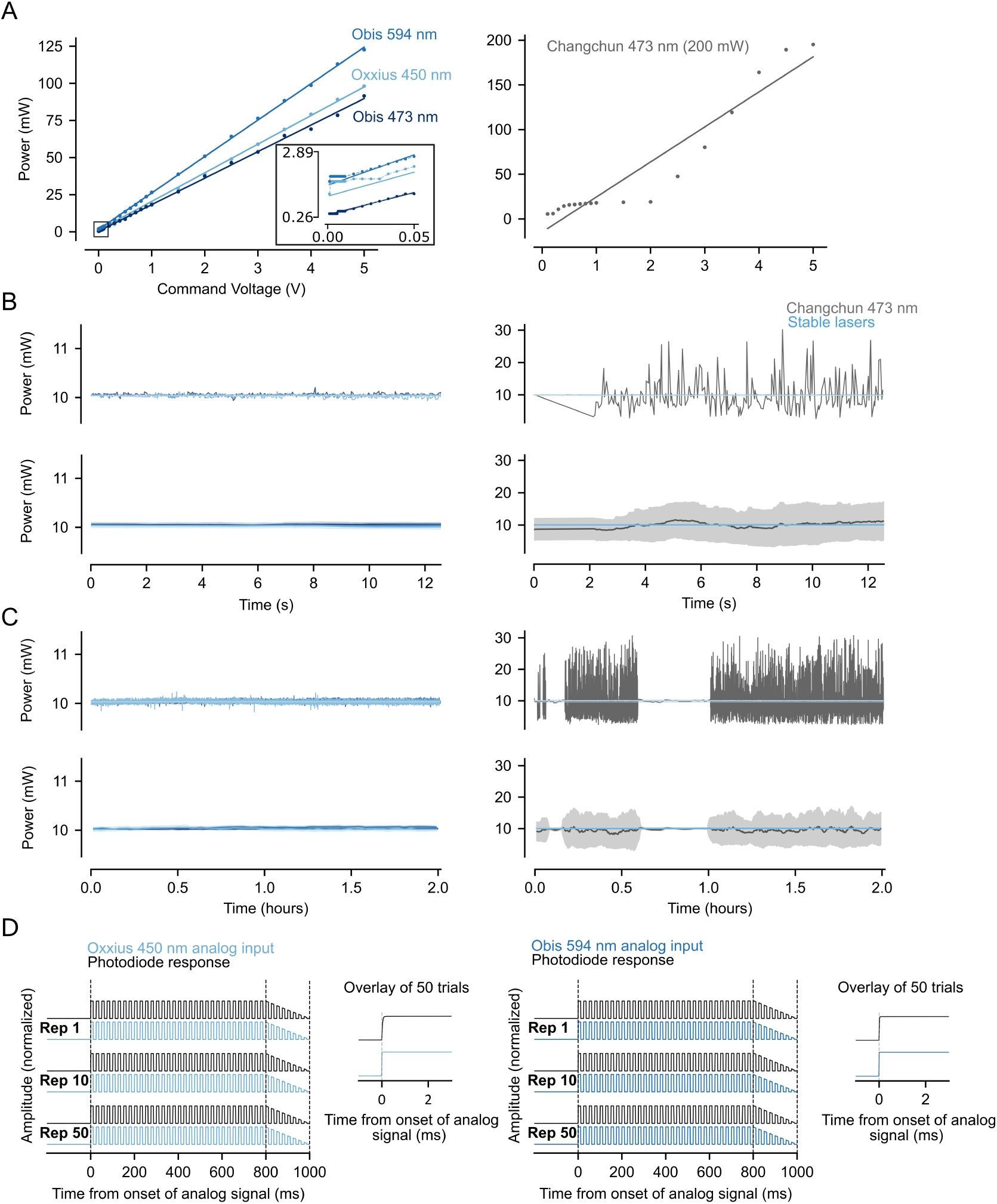
Laser tests for linearity, stability and onset responses. We tested Zapit with multiple lasers across a range of price points (3000 GBP - 10,000 GBP) with various wavelengths (450 nm to 594 nm). We also used a cheaper, less stable laser, for comparison (Changchun 473 nm). **A**. Power vs voltage plots for stable lasers (left, Obis 473 nm, Obis 594 nm, Oxxius 450 nm) and the unstable laser compared to stable lasers. Linear model fits were applied to the data. Inset: Zoom (black square) to show nonlinearity near 0 V, with linear model fit from the full data. **B**. Stability plots on short timescale for stable lasers (left) and all lasers (right). Top: raw power vs time. Bottom: drift analysis with a rolling average of data points within one window, each window 0.5 ms long from a 10 kHz sampling rate. Note the difference in y-axes between left and right. **C**. Stability plots on long timescale for stable lasers (left) and all lasers (right). Top: raw power vs time. Bottom: drift analysis with a rolling average of data points within one window, each window 10 minutes long from a 1 kHz sampling rate. **D**. Onset response plots for individual trials measuring latency of photodiode response relative to analog input for Oxxius 450 nm (left) and Obis 594 nm (right). Inset: 50 trials overlaid. Samples were acquired with a NI-DAQ USB-6453.

